# Analysis of cardiac magnetic resonance imaging traits in 29,000 individuals reveals shared genetic basis with dilated cardiomyopathy

**DOI:** 10.1101/2020.02.12.946038

**Authors:** James P. Pirruccello, Alexander Bick, Minxian Wang, Mark Chaffin, Steven A. Lubitz, Patrick T. Ellinor, Amit V Khera, Sekar Kathiresan, Krishna G. Aragam

**Affiliations:** Massachusetts General Hospital Division of Cardiology, Boston, Massachusetts; Center for Genomic Medicine, Massachusetts General Hospital, Boston, Massachusetts; Program in Medical and Population Genetics at the Broad Institute, Cambridge, Massachusetts; Harvard Medical School, Boston, Massachusetts; Verve Therapeutics, Cambridge, Massachusetts

## Abstract

Dilated cardiomyopathy (DCM) is an important cause of heart failure and the leading indication for heart transplantation. Many rare genetic variants have been associated with DCM, but common variant studies of the disease have yielded few associated loci. As structural changes in the heart are a defining feature of DCM, we conducted a genome-wide association study (GWAS) of cardiac magnetic resonance imaging (MRI)-derived left ventricular measurements in 29,041 UK Biobank participants. 26 novel loci were associated with cardiac structure and function. These loci were found near 17 genes previously shown to cause Mendelian cardiomyopathies. A polygenic score of left ventricular end systolic volume was associated with incident DCM in previously disease-free individuals (hazard ratio = 1.54 per one standard deviation increase in the polygenic score, P = 2.1×10^−16^). Even among carriers of truncating mutations in *TTN*, the polygenic score influenced the size and function of the heart. These results further implicate common genetic polymorphisms in DCM pathogenesis.

## Introduction

Affecting one in every 250 people, dilated cardiomyopathy (DCM) is a disease of cardiac muscle that leads to heart failure, and is the most common indication for cardiac transplantation^1^. Rare variants in dozens of genes have been associated with DCM ^2^. The most commonly identified mutations are truncating variants in *TTN* (TTNtv), which are found in 15-20% of DCM cases. However, rare variants in cardiomyopathy-related genes yield a genetic diagnosis for DCM in approximately 40% of cases^2^. Furthermore, sequencing efforts in large populations are now identifying an increasing number of individuals who harbor rare DCM-associated variants, but do not manifest clinical disease^3–6^.

A putative explanation for the limited diagnostic yield and incomplete penetrance of rare DCM-associated variants is that common genetic variation plays a role in the genetic architecture of DCM. To gain insight into the relationship between common genetic variants and DCM, one exome-wide association study^7^ and two genome-wide association studies (GWAS)^8,9^ have been conducted. Nine loci with variants that distinguished cases from controls at a genome-wide level of significance were identified, and four of these loci contain genes that also harbor rare DCM-causing mutations (*TTN, ALPK3, BAG3, FLNC*). These studies have successfully linked common variants to DCM, though their yield has been limited by modest sample sizes, as each study had fewer than 5,000 cases.

As structural changes to the left ventricle are a defining–and frequently incipient–feature of DCM, genetic analyses of cardiac imaging traits present another plausible method for genetic discovery^10,11^. Several GWAS of cardiac imaging phenotypes have probed the link between common genetic variants and changes in cardiac structure and function. For example, a large-scale genetic study of over 46,000 participants with transthoracic echocardiography (TTE)-derived phenotypes identified five genome-wide significant loci. Notably, one locus, near *PLN*, is associated with Mendelian cardiomyopathies^12,13^. A subsequent study using TTE in 19,000 participants from BioBank Japan yielded five additional novel loci, including a locus near *VCL* that is also associated with cardiomyopathies^14–16^. Finally, a study of 6,765 African American participants with cardiac imaging, including 1,210 with cardiac magnetic resonance imaging (MRI) from the Multi-ethnic Study of Atherosclerosis (MESA), yielded four suggestive loci that have not been subsequently confirmed^17^. Together, these studies highlight the potential value of population-based cardiac imaging to gain clinical and biological insights into myocardial diseases. Whether common genetic variants related to the heart’s structure and function can identify as-yet unaffected individuals at risk for DCM remains uncertain.

In this study, we sought to discover common genetic variants associated with cardiac structure and function, and to assess whether those common variants influenced the propensity of unrelated individuals to develop DCM. We performed these analyses in the UK Biobank, a population-based cohort of over 500,000 participants^18,19^. Within this cohort, a large imaging sub-study is underway, with plans to perform cardiac MRI in 100,000 participants^20,21^. The phenotypic characteristics of the first 5,000 participants have been described in detail^22,23^. Here, we analyzed automated measurements from the first 29,041 participants—the largest available cardiac MRI dataset—in order to study the relationship between common genetic variants, cardiac phenotypes, and risk for the development of DCM.

## Results

### Phenotype refinement and cardiac MRI results

We identified 29,041 UK Biobank participants with cardiac MRI readings who had no diagnosis of congestive heart failure (CHF), coronary artery disease (CAD), or DCM at the time of enrollment (**Supplemental Table 1** and **Supplemental Figure 1**). For these individuals, seven cardiac MRI-derived phenotypes were available: left ventricular end diastolic volume (LVEDV), left ventricular end systolic volume (LVESV), stroke volume (SV), the body-surface-area indexed versions of each of these traits (LVEDVi, LVESVi, and SVi), and left ventricular ejection fraction (LVEF). These individuals were majority female (52.6%) and had a mean age 63.6 years at the time of MRI. For women, the mean LVEDV, LVESV, and SV were 122.6mL, 41.1mL, and 81.5mL, respectively; for men, they were 152.8mL, 57.4mL, and 95.4mL (**Supplemental Table 2**). The LVEF averaged 0.67 for women and 0.63 for men (**Supplemental Figure 2**).

**Table 1:** Genome-wide significant loci from genetic association analyses of cardiac MRI phenotypes

| Locus ID | dbSNP | Chr | Position (hg19) | Effect Allele | Other Allele | Effect Allele Freq | INFO | Beta | SE | P-value | Locus Name |
| --- | --- | --- | --- | --- | --- | --- | --- | --- | --- | --- | --- |
| **LVEDV** |  |  |  |  |  |  |  |  |  |  |  |
| 1 | rs28579893 | 1 | 16347534 | A | G | 0.33 | 1.00 | -1.06 | 0.191 | 2.40E-08 | CLCNKA |
| 2 | rs4241246 | 2 | 71676366 | C | A | 0.43 | 0.99 | 1.04 | 0.184 | 1.10E-08 | DYSF |
| 3 | rs2562845 | 2 | 179514433 | T | C | 0.80 | 1.00 | 1.71 | 0.226 | 6.50E-14 | TTN |
| 4 |  | 3 | 14291003 | CTTTT | C | 0.66 | 0.98 | 1.13 | 0.192 | 4.20E-09 | TMEM43 |
| 5 |  | 3 | 99522473 | TA | T | 0.32 | 0.99 | -1.09 | 0.194 | 2.40E-08 | COL8A1 |
| 6 | rs1499813 | 3 | 171760427 | T | C | 0.59 | 1.00 | 0.99 | 0.184 | 4.20E-08 | FNDC3B |
| 7 | rs11153730 | 6 | 118667522 | T | C | 0.51 | 1.00 | -1.16 | 0.181 | 2.20E-10 | PLN |
| 8 |  | 8 | 125858538 | GA | G | 0.69 | 0.98 | 1.20 | 0.197 | 1.20E-09 | MTSS1 |
| 9 | rs72840788 | 10 | 121415685 | G | A | 0.78 | 0.98 | 1.67 | 0.222 | 4.80E-14 | BAG3 |
| 10 | rs3184504 | 12 | 111884608 | T | C | 0.47 | 1.00 | -1.22 | 0.181 | 1.10E-11 | SH2B3 |
| 11 |  | 15 | 85348961 | TTTTG | T | 0.75 | 0.97 | -1.16 | 0.211 | 3.00E-08 | ALPK3 |
| 12 | rs71385734 | 16 | 2160503 | T | G | 0.83 | 0.99 | 1.52 | 0.241 | 3.40E-10 | PKD1 |
| 13 | rs7502466 | 17 | 1372970 | G | A | 0.89 | 0.98 | 1.72 | 0.292 | 4.00E-09 | MYO1C |
| 14 | rs12460541 | 19 | 46312077 | G | A | 0.65 | 1.00 | -1.13 | 0.189 | 2.30E-09 | RSPH6A |
| **LVESV** |  |  |  |  |  |  |  |  |  |  |  |
| 15 | rs114300540 | 1 | 6248182 | C | T | 0.88 | 0.90 | -0.98 | 0.171 | 4.80E-09 | RPL22 |
| 1 | rs945425 | 1 | 16348412 | T | C | 0.32 | 1.00 | -1.06 | 0.114 | 6.30E-21 | CLCNKA |
| 16 | rs111323943 | 2 | 45610917 | C | CT | 0.96 | 0.90 | -1.46 | 0.270 | 4.00E-08 | SRBD1 |
| 2 | rs4852257 | 2 | 71678520 | T | G | 0.42 | 0.99 | 0.59 | 0.109 | 2.80E-08 | DYSF |
| 3 | rs2255167 | 2 | 179558282 | T | A | 0.81 | 1.00 | 1.30 | 0.136 | 2.30E-21 | TTN |
| 17 | rs1394785 | 2 | 213210974 | G | C | 0.29 | 0.99 | 0.65 | 0.118 | 4.10E-08 | ERBB4 |
| 4 | rs55691578 | 3 | 14277060 | A | C | 0.66 | 0.98 | 1.00 | 0.114 | 3.00E-18 | TMEM43 |
| 18 |  | 3 | 52890521 | AC | A | 0.15 | 0.99 | 0.81 | 0.150 | 1.30E-08 | TNNC1 |
| 6 | rs1499813 | 3 | 171760427 | T | C | 0.59 | 1.00 | 0.63 | 0.109 | 2.70E-09 | FNDC3B |
| 19 | rs34373805 | 7 | 128486363 | C | T | 0.84 | 1.00 | 0.81 | 0.147 | 1.30E-08 | FLNC |
| 8 |  | 8 | 125858538 | GA | G | 0.69 | 0.98 | 0.79 | 0.117 | 1.70E-11 | MTSS1 |
| 20 | rs1962104 | 8 | 141635329 | T | C | 0.45 | 0.98 | -0.62 | 0.109 | 1.90E-08 | AGO2 |
| 9 | rs72840788 | 10 | 121415685 | G | A | 0.78 | 0.98 | 1.52 | 0.132 | 7.10E-31 | BAG3 |
| 21 | rs113819537 | 12 | 26348429 | C | G | 0.75 | 0.99 | -0.71 | 0.124 | 7.50E-09 | SSPN |
| 22 | rs116904997 | 12 | 120668534 | G | A | 0.98 | 0.99 | -2.39 | 0.365 | 8.20E-11 | PXN |
| 23 | rs111418232 | 14 | 89835905 | A | T | 0.55 | 0.87 | -0.63 | 0.116 | 3.70E-08 | FOXN3 |
| 11 |  | 15 | 85348961 | TTTTG | T | 0.75 | 0.97 | -0.83 | 0.126 | 2.70E-11 | ALPK3 |
| 24 | rs5029142 | 16 | 988070 | T | A | 0.62 | 0.99 | 0.63 | 0.112 | 1.10E-08 | LMF1 |
| 12 | rs71385734 | 16 | 2160503 | T | G | 0.83 | 0.99 | 0.88 | 0.144 | 1.20E-09 | PKD1 |
| 25 | rs117837409 | 16 | 4631802 | A | G | 0.95 | 0.92 | -1.57 | 0.264 | 3.30E-09 | C16orf96 |
| 13 | rs7502466 | 17 | 1372970 | G | A | 0.89 | 0.98 | 1.07 | 0.174 | 1.20E-09 | MYO1C |
| 26 | rs10445344 | 17 | 53367788 | T | A | 0.73 | 0.98 | 0.68 | 0.122 | 1.10E-08 | HLF |
| 14 | rs10421891 | 19 | 46315809 | A | G | 0.64 | 1.00 | -0.75 | 0.112 | 7.40E-12 | RSPH6A |
| 27 | rs2267038 | 22 | 24169196 | G | C | 0.19 | 1.00 | -0.93 | 0.137 | 4.70E-12 | SMARCB1 |
| LVEF |  |  |  |  |  |  |  |  |  |  |  |
| 28 | rs2503715 | 1 | 2144107 | A | G | 0.13 | 0.91 | -0.00358 | 0.00067 | 4.10E-08 | C1orf86 |
| 1 | rs9442216 | 1 | 16353400 | T | C | 0.33 | 1.00 | 0.00428 | 0.00045 | 1.10E-21 | CLCNKA |
| 29 | rs10925197 | 1 | 236842077 | C | G | 0.46 | 0.99 | 0.00237 | 0.00043 | 2.50E-08 | ACTN2 |
| 3 | rs2255167 | 2 | 179558282 | T | A | 0.81 | 1.00 | -0.00423 | 0.00054 | 1.80E-15 | TTN |
| 4 | rs73028853 | 3 | 14273117 | C | T | 0.66 | 0.99 | -0.00382 | 0.00045 | 3.50E-17 | TMEM43 |
| 30 | rs62253176 | 3 | 69906231 | T | G | 0.81 | 0.99 | -0.00343 | 0.00055 | 8.70E-10 | MITF |
| 31 | rs4835730 | 5 | 138763807 | T | C | 0.33 | 1.00 | 0.00250 | 0.00046 | 4.10E-08 | DNAJC18 |
| 32 | rs3176326 | 6 | 36647289 | G | A | 0.80 | 0.99 | -0.00381 | 0.00054 | 1.80E-12 | CDKN1A |
| 19 | rs34373805 | 7 | 128486363 | C | T | 0.84 | 1.00 | -0.00355 | 0.00059 | 8.70E-10 | FLNC |
| 9 | rs72840788 | 10 | 121415685 | G | A | 0.78 | 0.98 | -0.00568 | 0.00053 | 5.60E-28 | BAG3 |
| 33 | rs58460019 | 10 | 129773144 | A | ATCAT<br>ACCT | 0.75 | 0.90 | 0.00285 | 0.00052 | 2.20E-08 | PTPRE |
| 11 | rs8023658 | 15 | 85323220 | G | T | 0.51 | 0.96 | -0.00275 | 0.00044 | 4.40E-10 | ALPK3 |
| 24 | rs5015439 | 16 | 987358 | T | C | 0.62 | 1.00 | -0.00242 | 0.00044 | 1.80E-08 | LMF1 |
| 26 | rs3794752 | 17 | 53382829 | T | C | 0.72 | 0.98 | -0.00286 | 0.00048 | 6.30E-10 | HLF |
| 34 | rs2047273 | 18 | 34184859 | T | C | 0.68 | 0.96 | -0.00256 | 0.00047 | 3.10E-08 | FHOD3 |
| 27 | rs2070458 | 22 | 24159307 | A | T | 0.20 | 1.00 | 0.00361 | 0.00054 | 1.40E-11 | SMARCB1 |
| **SV** |  |  |  |  |  |  |  |  |  |  |  |
| 3 | rs7573293 | 2 | 179753245 | C | T | 0.28 | 0.99 | -0.699 | 0.124 | 1.60E-08 | TTN |
| 7 | rs3951016 | 6 | 118559658 | T | A | 0.53 | 0.99 | 0.692 | 0.111 | 5.20E-10 | PLN |
| 10 | rs11065979 | 12 | 112059557 | C | T | 0.57 | 1.00 | 0.714 | 0.112 | 1.60E-10 | BRAP |
| **LVEDVi** |  |  |  |  |  |  |  |  |  |  |  |
| 3 | rs2042995 | 2 | 179558366 | T | C | 0.77 | 1.00 | 0.808 | 0.107 | 5.20E-14 | TTN |
| 4 | rs2170454 | 3 | 14294549 | T | C | 0.46 | 0.99 | -0.505 | 0.091 | 4.10E-08 | TMEM43 |
| 7 | rs72967533 | 6 | 118655020 | T | C | 0.52 | 0.99 | -0.529 | 0.091 | 7.90E-09 | PLN |
| 8 |  | 8 | 125858538 | GA | G | 0.69 | 0.98 | 0.546 | 0.098 | 2.00E-08 | MTSS1 |
| 9 | rs11199043 | 10 | 121367249 | A | G | 0.52 | 0.96 | -0.620 | 0.092 | 1.80E-11 | BAG3 |
| 10 | rs653178 | 12 | 112007756 | C | T | 0.47 | 1.00 | -0.549 | 0.090 | 1.00E-09 | ATXN2 |
| 13 | rs7502466 | 17 | 1372970 | G | A | 0.89 | 0.98 | 0.817 | 0.146 | 1.80E-08 | MYO1C |
| 14 | rs12460541 | 19 | 46312077 | G | A | 0.65 | 1.00 | -0.544 | 0.095 | 1.00E-08 | RSPH6A |
| **LVESVi** |  |  |  |  |  |  |  |  |  |  |  |
| 28 | rs2503715 | 1 | 2144107 | A | G | 0.13 | 0.91 | 0.447 | 0.081 | 2.40E-08 | C1orf86 |
| 1 | rs9442216 | 1 | 16353400 | T | C | 0.33 | 1.00 | -0.471 | 0.054 | 5.70E-19 | CLCNKA |
| 3 | rs2042995 | 2 | 179558366 | T | C | 0.77 | 1.00 | 0.574 | 0.061 | 4.50E-21 | TTN |
| 17 | rs4673661 | 2 | 213205047 | A | G | 0.28 | 1.00 | 0.322 | 0.057 | 7.60E-09 | ERBB4 |
| 4 | rs73028849 | 3 | 14272766 | G | C | 0.66 | 0.99 | 0.455 | 0.054 | 9.10E-17 | TMEM43 |
| 6 | rs1499813 | 3 | 171760427 | T | C | 0.59 | 1.00 | 0.289 | 0.052 | 9.40E-09 | FNDC3B |
| 32 | rs3176326 | 6 | 36647289 | G | A | 0.80 | 0.99 | 0.383 | 0.065 | 4.30E-09 | CDKN1A |
| 19 | rs34373805 | 7 | 128486363 | C | T | 0.84 | 1.00 | 0.380 | 0.071 | 3.40E-08 | FLNC |
| 8 |  | 8 | 125858538 | GA | G | 0.69 | 0.98 | 0.342 | 0.056 | 6.70E-10 | MTSS1 |
| 20 | rs1962104 | 8 | 141635329 | T | C | 0.45 | 0.98 | -0.316 | 0.052 | 2.20E-09 | AGO2 |
| 9 | rs72840788 | 10 | 121415685 | G | A | 0.78 | 0.98 | 0.671 | 0.063 | 3.70E-26 | BAG3 |
| 21 | rs113819537 | 12 | 26348429 | C | G | 0.75 | 0.99 | -0.324 | 0.059 | 3.50E-08 | SSPN |
| 22 | rs116904997 | 12 | 120668534 | G | A | 0.98 | 0.99 | -1.129 | 0.175 | 4.30E-10 | PXN |
| 11 |  | 15 | 85348961 | TTTTG | T | 0.75 | 0.97 | -0.347 | 0.060 | 7.40E-09 | ALPK3 |
| 24 | rs5029140 | 16 | 987972 | C | T | 0.62 | 0.99 | 0.320 | 0.053 | 3.90E-10 | LMF1 |
| 13 | rs7502466 | 17 | 1372970 | G | A | 0.89 | 0.98 | 0.475 | 0.083 | 1.50E-08 | MYO1C |
| 26 | rs876719 | 17 | 53366566 | G | T | 0.73 | 0.97 | 0.354 | 0.059 | 7.10E-10 | HLF |
| 14 | rs10421891 | 19 | 46315809 | A | G | 0.64 | 1.00 | -0.336 | 0.054 | 1.90E-10 | RSPH6A |
| 27 | rs2070458 | 22 | 24159307 | A | T | 0.20 | 1.00 | -0.460 | 0.065 | 5.20E-13 | SMARCB1 |
| **SVi** |  |  |  |  |  |  |  |  |  |  |  |
| 2 | rs7573293 | 2 | 179753245 | C | T | 0.28 | 0.99 | -0.365 | 0.063 | 1.00E-08 | TTN |
| 35 |  | 2 | 232288831 | TTTC | T | 0.68 | 0.99 | -0.371 | 0.061 | 1.50E-09 | B3GNT7 |
| 36 | rs2146324 | 6 | 43756863 | A | C | 0.26 | 0.98 | 0.357 | 0.065 | 4.30E-08 | VEGFA |
| 7 | rs3951016 | 6 | 118559658 | T | A | 0.53 | 0.99 | 0.318 | 0.057 | 2.80E-08 | PLN |
| 10 | rs11065979 | 12 | 112059557 | C | T | 0.57 | 1.00 | 0.321 | 0.057 | 2.40E-08 | BRAP |
| 37 | rs7143356 | 14 | 23881083 | T | C | 0.62 | 1.00 | 0.319 | 0.058 | 4.10E-08 | MYH7 |

**Figure 1:**
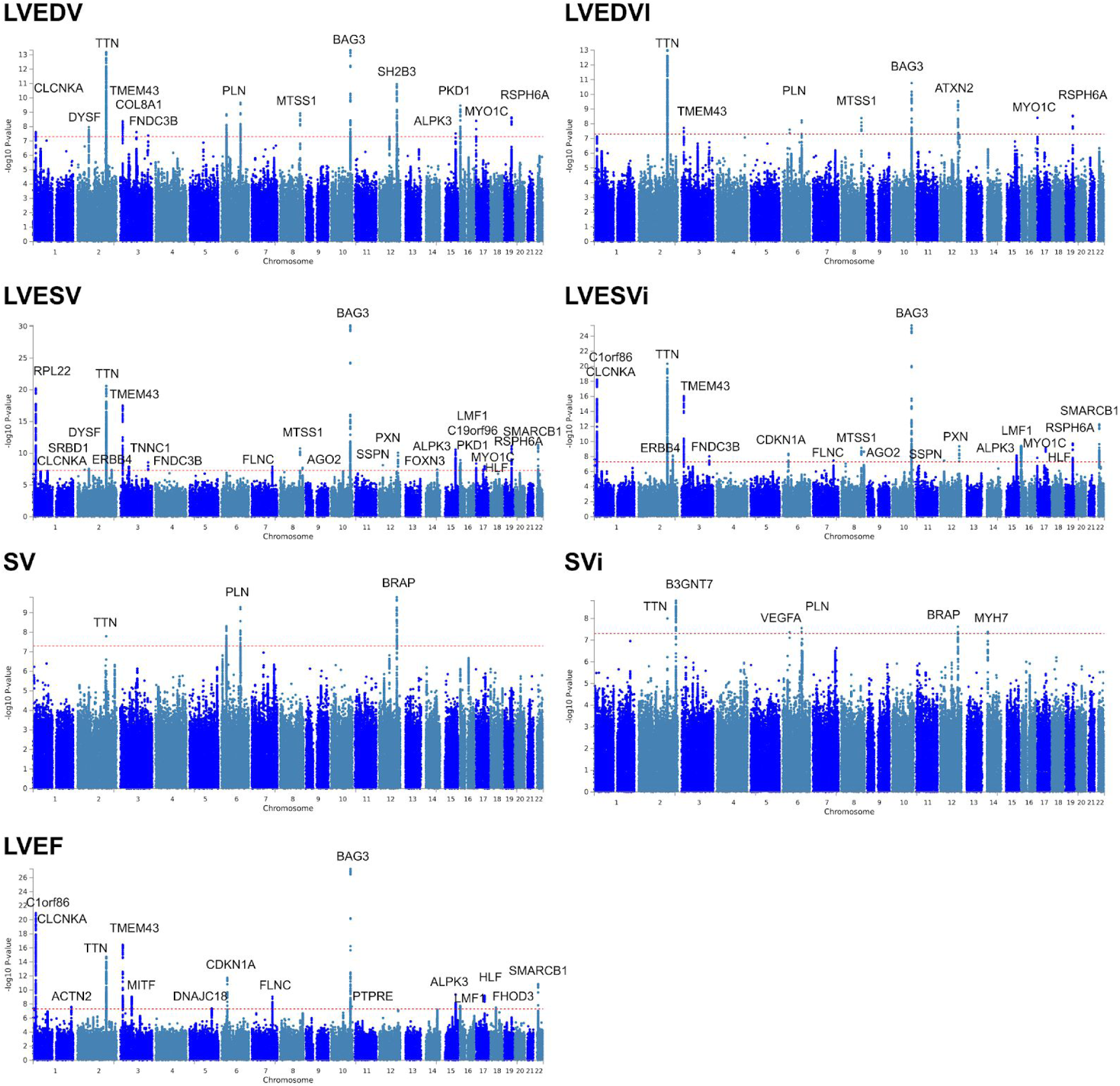
Manhattan plots of genome-wide association discovery analyses of cardiac MRI phenotypes in the UK Biobank. For each cardiac MRI phenotype, the -log10(P value) is graphed on the y-axis at each chromosomal position on the x-axis. The nearest gene to each genome-wide significant lead SNP is labeled at each locus, except when a cardiomyopathy-related gene is present within 250kb of the lead SNP.

**Figure 2:**
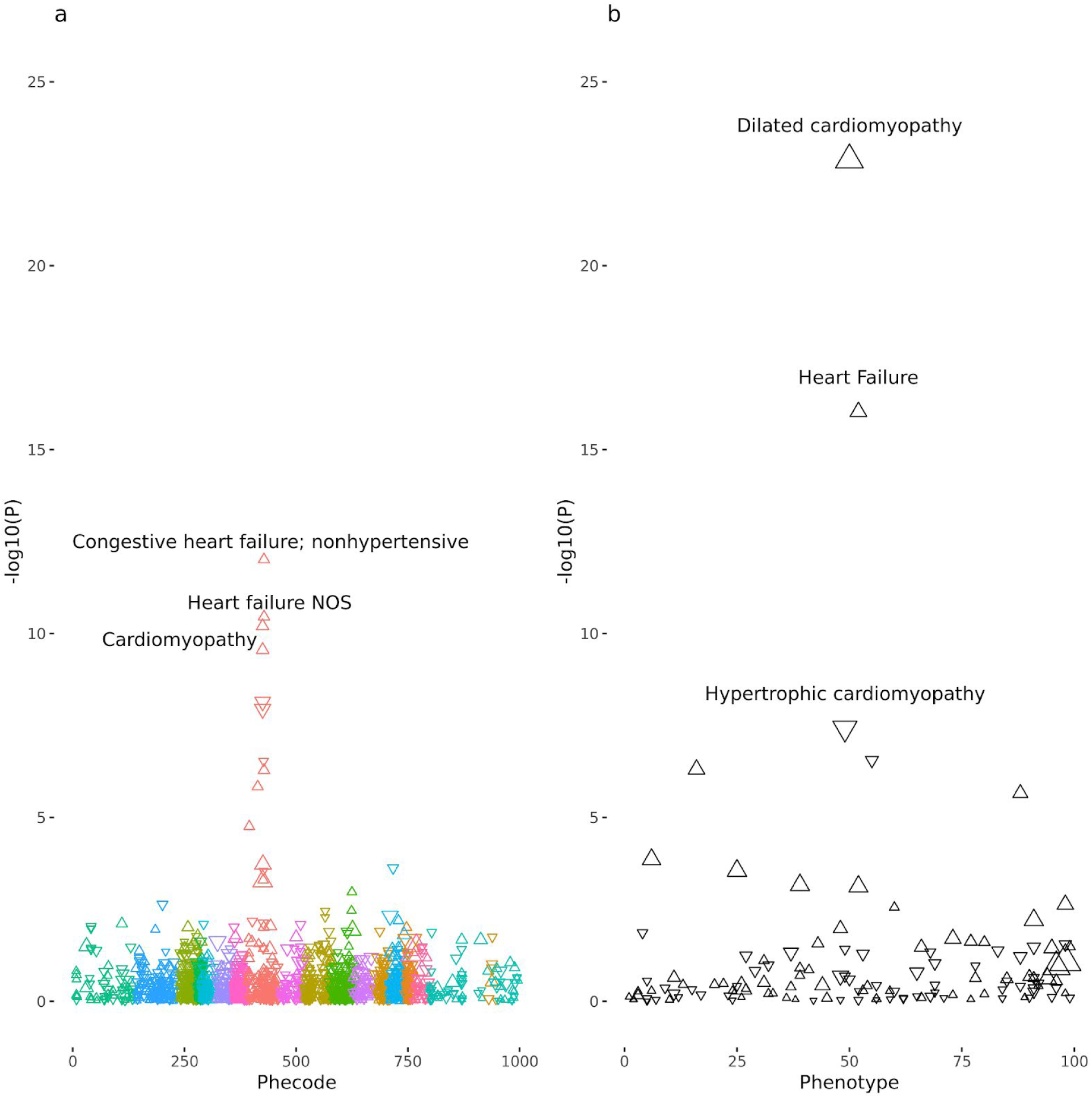
PheWAS highlights the connection between a polygenic score for LVESVi and dilated cardiomyopathy. The polygenic score derived from LVESVi was applied to PheCodes (panel a) and curated disease phenotypes (panel b) in the UK Biobank. Each of the curated phenotypes is defined in **Supplemental Table 1**. For both panels (a) and (b), the X-axis represents the identifying code for the disease phenotype. The Y-axis represents the -log10 of the P-value of association between the polygenic score and the phenotype in a logistic model adjusted for age at enrollment, the genotyping array, sex, and the first five principal components of ancestry. Triangles oriented upward represent betas that are concordant with the LVESVi PRS (e.g., a higher LVESVi PRS corresponds with a higher risk of DCM), and the reverse is true for downward-oriented triangles. The 3 most strongly associated phenotypes in each panel are labeled on the figure. The PheWAS plots for all seven cardiac MRI phenotypes are available in **Supplemental Figure 7**.

### Cardiac structure and function are heritable

We asked whether the cardiac MRI phenotypes were influenced by participants’ genetic backgrounds. SNP-based heritability—the proportion of variance explained by all SNPs on a genotyping array—was 45% for LVEDV, 43% for LVESV, and 31% for LVEF. Genetic correlations between these cardiac MRI phenotypes were similar in magnitude to their epidemiologic correlations (**Supplemental Figures 3** and **4**).

**Figure 3:**
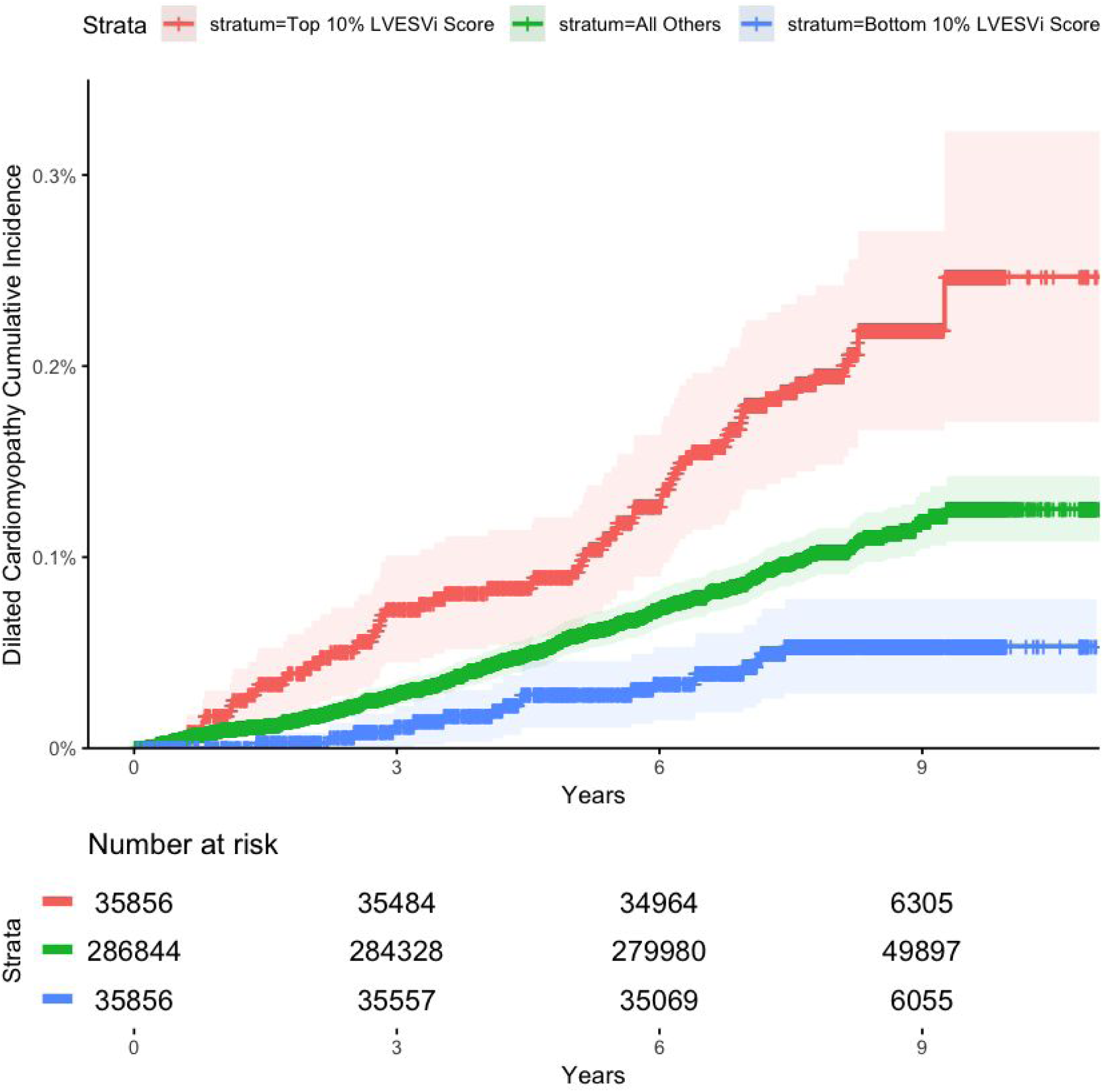
The LVESVi polygenic score influences risk for incident dilated cardiomyopathy. The cumulative DCM incidence over time is plotted for individuals in the bottom tenth percentile (blue), middle 80% (green) and top tenth percentile (red) for the LVESVi polygenic score. The 95% confidence intervals are represented with lighter colors. The X-axis represents the number of years since enrollment in the UK Biobank. The Y axis represents the cumulative incidence of DCM.

### 37 genetic loci are associated with left ventricular size and function

Having established a genetic basis for variability in cardiac structure and function, we then performed a GWAS to identify common genetic variants associated with the seven cardiac MRI phenotypes. 37 distinct loci in the human genome were associated with one or more cardiac MRI phenotypes at genome-wide significance (P < 5×10^−8^; **Table 1**). LD score regression revealed modest test statistic inflation, consistent with polygenicity rather than population stratification (**Supplemental Table 3**).

Of the 37 genome-wide significant loci, 26 were novel and had not been described in prior common variant analyses of dilated cardiomyopathy or cardiac imaging phenotypes. In total, 14 loci associated with LVEDV, eight with LVEDVi, 24 with LVESV, 19 with LVESVi, 16 with LVEF, three with SV, and six with SVi (**Figure 1**).

Among our genome-wide significant loci, we found five of the 10 loci previously discovered through genetic analyses of echocardiographic measurements, including SNPs near *PLN, SH2B3/ATXN2, MTSS1, SMARCB1*, and *CDKN1A*^13,14^. In addition, we recovered six of the nine loci previously identified at exome- or genome-wide significance in common variant studies of heart failure or cardiomyopathy, including SNPs near *ALPK3, BAG3, CLCNKA/HSPB7, FHOD3, FLNC*, and *TTN*^7–9^.

### GWAS loci are enriched for genes expressed in cardiac and skeletal muscle

In order to understand the tissue specificity of the discovered loci, we applied MAGMA, which revealed an enrichment of variants clustered near genes expressed in cardiac and skeletal muscle tissue types, with results for LVESV, LVESVi, and LVEF achieving a Bonferroni-corrected significance threshold (**Supplemental Figure 5**).

### GWAS loci are enriched for Mendelian cardiomyopathy-linked genes

We then asked whether the 37 GWAS loci were in proximity to more Mendelian cardiomyopathy genes (from the panel of 129 genes in **Supplemental Table 4**) than expected based on chance. Out of 100,000 simulations, none was linked to more than 14 Mendelian genes (**Supplemental Figure 6a**); in contrast, our loci are near 17 Mendelian genes from the gene panel (**Supplemental Table 5**), were significantly enriched for Mendelian cardiomyopathy genes (P < 1×10^−5^). For comparison, the prior cardiomyopathy GWAS studies identified loci near eight Mendelian cardiomyopathy genes, of which seven are also identified in this study (**Supplemental Figure 6b**).

### PheWAS links cardiac MRI polygenic scores with dilated cardiomyopathy in a cross-sectional analysis

We also produced polygenic scores for each trait, weighting the genetic dosage by the effect size of the lead SNPs from each GWAS from **Table 1**. We performed a cross-sectional phenome-wide association study (PheWAS) in over 450,000 UK Biobank participants to assess the relationship between each of the seven polygenic scores and disease phenotypes. We first performed a PheWAS using a broad set of PheCodes^24^, 1,598 of which were present in 20 or more participants. As anticipated, this analysis showed an enrichment for cardiac diseases. We then performed a PheWAS for 96 diseases using curated definitions (as defined in **Supplemental Table 1**). Among our curated disease traits, DCM emerged as the disease most strongly associated with the polygenic scores for LVEF, LVEDV, LVESV, LVESVi. The LVESVi polygenic score had the single strongest relationship with DCM (OR 1.47 per standard deviation [SD] increase in LVESVi polygenic score; P = 1.3×10^−23^; **Figure 2**). The strongest relationships between each of the 7 polygenic scores and the disease phenotypes are shown in **Supplemental Table 6** and **Supplemental Figure 7**.

### A polygenic score for LVESVi is associated with incident dilated cardiomyopathy

Having established a strong relationship between the common variant-derived polygenic scores identified in our study and DCM in the cross-sectional PheWAS, we then asked whether these polygenic scores also predicted incident disease. The association between polygenic scores and incident DCM (388 cases) was assessed in the remaining 358,556 individuals in the UK Biobank without cardiac MRI data, after excluding those with cardiac disease at baseline, and those with third-degree or closer relatedness to another participant (**Supplemental Table 7**). The polygenic scores for LVEDV, LVESV, LVEF, LVEDVi, and LVESVi were significantly associated with incident DCM after adjusting for age, sex, genotyping batch, and the first five principal components of ancestry in separate Cox proportional hazards models for each phenotype.

The 19-SNP LVESVi polygenic score had the strongest relationship with incident DCM (hazard ratio [HR] = 1.54 per SD increase in the score, P = 2.1×10^−16^). The direction was consistent with clinical expectations: a greater genetically mediated LVESVi corresponded with a higher risk of DCM, while a lower polygenic risk corresponded to a lower risk of DCM. The cumulative incidence of DCM among those in the top 10%, bottom 10%, and middle 80% for the polygenic score are plotted in **Figure 3**. For each trait, the relationship between its polygenic score and incident DCM is available in **Supplemental Table 8**.

### A polygenic score for LVESVi influences left ventricular structure and function in individuals harboring rare truncating variants in *TTN*

Finally, to understand whether common genetic variants might contribute to the incomplete penetrance and variable expressivity of DCM-associated rare variants, we asked whether the 19-SNP LVESVi polygenic score affected the structure and function of the heart among carriers of high-impact truncating mutations in *TTN* (TTNtv). Rare TTNtv were chosen for analysis because of their established roles in cardiomyopathy and preclinical cardiac dysfunction, as well as their relatively high population frequency (approximately 0.5%). Among the 11,878 participants who had undergone both cardiac MRI and exome sequencing, we identified 53 carriers of TTNtv in exons spliced into over 90% of transcripts in the heart (**Supplemental Table 9**). We found that the polygenic background of TTNtv carriers influenced both left ventricular volume and function: the LVEDV of TTNtv carriers was influenced by the LVESVi polygenic score (12.0mL increase in LVEDV per SD increase in the score, P = 0.011), as was LVESV (9.1mL increase in LVESV per SD, P = 0.008). The LVEF was also reduced by 2.8% per SD of the LVESVi score (P = 0.017).

## Discussion

In this study, we identified 26 novel, common genetic variants associated with the structure and function of the heart in individuals without overt cardiovascular disease; linked our identified common variants to DCM through multiple lines of evidence; established that a polygenic score derived from common genetic variants predicts incident DCM; and demonstrated that a common genetic background affects cardiac traits even among carriers of cardiomyopathy-related rare variants.

These results permit several conclusions. First, common genetic variants associated with left ventricular structure and function contribute to DCM risk. In an incident disease analysis, our best performing polygenic score—created from 19 SNPs from our LVESVi GWAS—robustly predicted DCM (HR 1.54 per standard deviation increase, P = 2 × 10^−16^). This is the first instrument derived from common genetic variants that has been demonstrated to associate with incident DCM. That the common genetic variants contributing to the LVESVi polygenic score were identified after excluding individuals with any known cardiac disease—yet still predicted cardiomyopathy—suggests an intrinsic connection between the common genetic determinants of normal cardiac traits and risk for DCM. These results raise the possibility that, in some individuals, DCM may reflect the extremes of normal phenotypic variation, driven by a high burden of common variants. This may, in part, explain the incomplete yield of genetic testing for rare variants in DCM. Future studies are required to determine the relative contributions of—and potential interplay between—common variants, rare variants, and environmental factors in the pathogenesis of DCM.

Second, genetic analyses of quantitative cardiac imaging traits may improve our understanding of the common genetic basis of cardiomyopathies. In our prior genetic analysis of the UK Biobank, we refined a heterogeneous heart failure phenotype to a specific, nonischemic cardiomyopathy subset, enabling detection of two DCM risk loci (near *BAG3* and *CLCNKA*) that associated with subclinical changes in LV structure and function^25^. Similarly, seminal GWAS studies of dilated cardiomyopathy yielded 9 risk loci, but were limited by the recruitment of cardiomyopathy cases^7–9^. Here, by pursuing a genetic analysis of cardiac MRI-derived traits in individuals without clinical disease, we recovered the majority of previously reported cardiomyopathy risk loci, and discovered loci near an additional 10 Mendelian cardiomyopathy genes (**Supplemental Figure 6**). Additionally, our PheWAS confirmed in an unbiased fashion that the genetic loci from these phenotypes were strongly associated with cardiomyopathies. Further analyses of MRI-derived cardiac traits may permit efficient study of the genetic determinants of cardiac structure and function in both health and disease, complementing the growing case-control genetic studies of cardiomyopathies.

Third, our results provide new insights into the role that common genetic variants play in determining the structure and function of the heart, even in the context of rare, high-impact mutations in cardiomyopathy-related genes. In a prior analysis of TTNtv, TTNtv carrier status associated with changes in LVEDV (+11.8mL), LVESV (+7.7mL), and LVEF (−2.8%) in individuals without clinical DCM^6^. In the present study, among carriers of cardiac-relevant TTNtv (those spliced into at least 90% of transcripts in the heart), a 1-SD increase in the LVESVi polygenic score associated with comparable changes in LVEDV (+12.0mL), LVESV (+9.1mL), and LVEF (−2.8%). These findings emphasize the potential scope and impact of common genetic variants on cardiac structure and function, and suggest that the penetrance of high-impact rare variants may be influenced by carriers’ polygenic backgrounds: for example, individuals with a TTN truncation but a favorable genetic background may be less likely to develop a reduced LVEF. Future studies will be required to understand if these individuals might be protected from cardiomyopathy.

Finally, our results highlight the importance of costameres to variation in cardiac traits. Costameres are cytoskeletal assemblies that connect the sarcomere—the basic structural unit of muscle—to the cell membrane and the extracellular matrix. Costameres include protein complexes such as the dystrophin glycoprotein complex (DGC) and the vinculin-talin-integrin system^26^. Both the DGC and the vinculin-talin-integrin system play roles in mechanical anchoring, while the vinculin-talin-integrin system also plays a role in mechanosensation and signal transduction^27^. Notably, three lead SNPs’ nearest genes (*FLNC, SSPN*, and *PXN*) play key roles in costamere biology. Ɣ-filamin (produced by *FLNC*) links the membrane-bound DGC with sarcomeric actin filaments^28^. *PXN* produces a protein (paxillin) that binds to vinculin and integrin to form part of the vinculin-talin-integrin system^29^. Finally, sarcospan, the protein product of *SSPN*, is a part of the DGC and is required for DGC-integrin interactions^30,31^. The emergence of these loci from our unbiased genetic analysis of cardiac MRI phenotypes underscores the importance of the costamere within the cardiomyocyte, and suggests that compromise of its protein assemblies may contribute to the development of cardiomyopathies.

### Study limitations

There are several limitations to our findings. First, our analyses were limited to older individuals of predominantly European ancestry, which may limit their applicability to younger individuals and those of other ancestries. Second, the cardiac measurements are derived from automated readings. Although we excluded gross anatomical mistracings in images at the tails of the distribution, these exclusions would not correct for algorithmic biases in the automated reading software. Third, because the UK Biobank relied on hospitalization or death to assign disease status, unrecognized disease at baseline may have occurred for individuals without any pre-enrollment hospitalizations.

### Conclusions

In conclusion, we report 26 novel, common genetic loci associated with left ventricular structure and function as measured by cardiac MRI, and reveal a robust link to dilated cardiomyopathy.

## Methods

### Study participants

The UK Biobank is a richly phenotyped, prospective, population-based cohort ^19^. In total, we analyzed 487,283 participants with genetic data who had not withdrawn consent as of October 2018. Analysis of the UK Biobank data was approved by the Partners Health Care institutional review board (protocol 2013P001840).

### Phenotype refinement

Disease phenotypes in the UK Biobank were defined using a combination of self-reported data (confirmed by a healthcare professional), hospital admission data, and death registry data. We defined nonischemic dilated cardiomyopathy (DCM) as a billing code diagnosis of dilated cardiomyopathy (ICD-10 code I42.0) in the absence of coronary artery disease. Algorithms for identifying individuals with DCM, heart failure from any cause, and coronary artery disease in the UK Biobank are available in **Supplemental Table 1**, as previously described ^25^.

### Cardiac MRI measurements

Cardiac magnetic resonance imaging was performed with 1.5 Tesla scanners (MAGNETOM Aera, Syngo Platform vD13A, Siemens Healthcare) with electrocardiographic gating for cardiac synchronization ^21^. Cardiac assessment was performed from the combination of several cine series using balanced steady-state free precession acquisitions, with post-processing by cvi42 Version 5.1.1^22^. Because of known bias in the vD13A automated measurements, a bias correction was applied for left ventricular end diastolic volume (LVEDV) and left ventricular end systolic volume (LVESV) measurements, using linear corrections derived from a UK cohort undergoing imaging on the same MRI platform ^32^.

Values for left ventricular ejection fraction (LVEF) and stroke volume (SV) were calculated from the LVEDV and LVESV: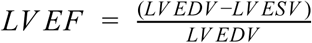 and *SV* = *LV EDV* − *LV ESV*. Body surface area (BSA)-indexed values were produced for left ventricular end diastolic volume (LVEDVi), left ventricular systolic volume (LVESVi), and stroke volume (SVi) by, respectively, dividing the values for LVEDV, LVESV, and SV by the individual’s Mosteller BSA: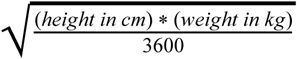^33^.

### Cardiac MRI sample selection and quality control

At the time of analysis in June 2019, we identified 31,931 individuals with cardiac MRI data. To account for errors in the automated volume measurement system, two cardiologists (JPP and KGA) manually reviewed images for samples having LVEDV or LVESV beyond 1.5 interquartile range below the first quartile or above the third quartile^34^. 610 samples identified as having gross anatomic mistracings were rejected. 1,308 samples that did not pass genotyping quality control, detailed below, were excluded. Data from 870 individuals with incident or prevalent heart failure, dilated cardiomyopathy, hypertrophic cardiomyopathy, or coronary artery disease were excluded. Finally, we removed 102 samples with LVEDV or LVESV beyond 4 standard deviations above or below the mean. After these exclusions, 29,041 samples remained for analysis (flow diagram displayed in **Supplemental Figure 1**).

### Genotyping and quality control

UK Biobank samples were genotyped on either the UK BiLEVE or UK Biobank Axiom arrays, then imputed into the Haplotype Reference Consortium panel and the UK10K + 1000 Genomes panel^18^.

We performed genotyping quality control by excluding genotyped variants with call rate < 0.95, imputed variants with INFO score < 0.3, imputed or hard-called variants with minor allele frequency less than 0.001 in the overall UK Biobank population, or imputed or hard-called variants with an effective minor allele count of less than 100 in the subset of individuals with cardiac MRI data. To calculate the effective minor allele count, the minor allele frequency was multiplied by the number of alleles in the analysis and by the imputation INFO score provided by the UK Biobank^18^. After exclusions, 13,660,711 autosomal variants were analyzed. Variant positions were keyed to GRCh37/hg19.

We performed sample quality control by excluding samples that had no imputed genetic data; a genotyping call rate < 0.98; a mismatch between submitted and inferred sex; sex chromosome aneuploidy (neither XX nor XY); exclusion from kinship inference; excessive third-degree relatives; or that were outliers in heterozygosity or genotype missingness rates as defined centrally by the UK Biobank^18^.

### Heritability analysis

Heritability attributable to single nucleotide polymorphisms (SNPs) at the 784,256 directly genotyped sites that passed quality control was computed with the --reml option from BOLT-LMM (version 2.3.2)^35^. Cross-trait genetic correlations between LVESV, LVEDV, and LVEF were computed with the same command.

### Genome-wide association study

We performed genome-wide association studies (GWAS) using linear mixed models with BOLT-LMM (version 2.3.2) to account for ancestral heterogeneity, cryptic population structure, and sample relatedness^35,36^.

Because the cardiac MRI data released by the UK Biobank comprised the majority of cardiac MRIs ever released in a research setting, we implemented a within-cohort replication strategy. First, we used the initial 24,216 samples with MRI data released by the UK Biobank through December 2018 to conduct a genome-wide association study (GWAS) for each of 7 cardiac MRI phenotypes: LVEDV, LVESV, LVEF, SV, LVEDVi, LVESVi, and SVi. Each GWAS was adjusted for the first five principal components of ancestry, sex, year of birth, age at the time of MRI, and the MRI scanner’s unique identifier to account for batch effects. We repeated each GWAS in the replication set of 4,825 samples that were made available in March 2019. With this replication strategy, we took SNPs forward for a final joint analysis if they (1) achieved P < 1×10^−4^ in the initial analysis and (2) had the same direction of effect in the replication analysis.

We then conducted a joint GWAS in all 29,041 samples for any SNP that achieved nominal replication. Variants with association P < 5×10^−8^ in the joint analysis were considered to be genome-wide significant. The most strongly associated SNP at each locus is referred to as the “lead SNP.” The gene in closest physical proximity to each lead SNP was designated as the gene label associated with each locus; however, if a Mendelian cardiac disease-associated gene was located within 250kb of the lead SNP, then the locus was named after the nearest Mendelian gene.

LD score regression was performed with ldsc (version 1.0.0) to partition the genomic control factor lambda into polygenic and inflation components^37^. Allelic heterogeneity was assessed by clumping at each genome-wide significant locus to identify additional variants having trait association P<5×10^−8^ and r^2^<0.2 with other independent genome-wide significant variants within 1 megabase of the top variant at each locus using FUMA^38^.

### Tissue enrichment

From the associated SNPs for each genotype, we evaluated the associations of 10,678 gene sets from MSigDB v6.2 and gene expression sets from GTEx using MAGMA^39–41^. Tissue enrichment tests were executed on the FUMA platform v1.3.4b^38^. A Bonferroni-corrected threshold of 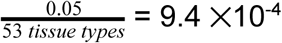 was considered statistically significant enrichment of a tissue type.

### Mendelian cardiomyopathy gene set enrichment

To create an unbiased list of genes associated with Mendelian cardiomyopathies and congenital heart abnormalities, we assembled genes from three commercially available cardiomyopathy gene panels (GeneDx Cardiomyopathy Panel, 207 Perry Parkway, Gaithersburg, MD, USA; Invitae Cardiomyopathy Comprehensive Panel, 1400 16th Street, San Francisco, CA, USA; Partners Laboratory for Molecular Medicine Pan Cardiomyopathy Panel, 77 Avenue Louis Pasteur Suite 250, Boston, MA, USA). Together, these panels contained 129 genes (**Supplemental Table 4**), all of which mapped to transcripts with HG19 coordinates. SNPsnap was used to generate 100,000 sets of SNPs that match the lead SNPs from the GWAS based on minor allele frequency, number of SNPs in linkage disequilibrium, distance to the nearest gene, and gene density at the locus^42^. A SNP was considered to be near a Mendelian locus if it was within 1 megabase upstream or downstream of any gene on the panel. Significance was assessed by permutation testing across the 100,000 SNP sets to determine the neutral expectation for the number of overlapping genes in loci with characteristics similar to ours.

### Polygenic score creation

For each individual, we calculated a polygenic score by taking the beta for each lead SNP multiplied by the genetic dosage of the effect allele, and summing this value for each lead SNP from **Table 1** for that trait. We repeated this procedure for each of the 7 traits, producing 7 polygenic scores.

### Phenome-wide association study

We performed a phenome-wide association study (PheWAS) in the 456,237 individuals with genetic data who had not undergone cardiac MRI. We tested polygenic scores produced for each of the 7 cardiac traits for association with 96 manually curated disease phenotypes and 1,598 ICD-10 mapped PheCodes which were present in 20 or more individuals in the UK Biobank^24^. Associations between the polygenic score and each phenotype were modeled with logistic regression, accounting for age at enrollment, sex, the genotyping array, and the first 5 principal components of ancestry as covariates.

### Testing polygenic score association with incident dilated cardiomyopathy

We assessed the relationship between the polygenic scores and DCM in the 358,556 participants who had not undergone cardiac MRI, were free from CHF, DCM, and CAD at baseline, and who were not identified by the UK Biobank as having third-degree or closer relatedness to another participant^18^. This sample included 388 individuals with incident DCM. We tested each score separately for association with incident DCM using a Cox proportional hazards model. This model was adjusted for sex, genotyping array, the first five principal components of ancestry, and the cubic basis spline of age at enrollment. Cox modeling was performed with the *survival* package in R 3.4.4 (R Foundation for Statistical Computing, Vienna, Austria). We defined statistical significance as a two-tailed P < (0.05 / 7 phenotypes) = 0.007.

For plotting, the samples were divided into deciles by LVESVi polygenic score. DCM incidence over time was plotted for the individuals in the lowest 10% of the LVESVi polygenic score, those in the highest 10%, and those in the remaining 80% using the *survminer* package in R.

### Testing the polygenic score within TTNtv carriers

As described by Van Hout, *et al*, samples from the UK Biobank were prioritized for exome sequencing based on the presence of MRI data and linked records^4^. Exomes were generated by the Regeneron Genetics Center, and reprocessed by the UK Biobank according to Functional Equivalent standards^43^. In total, 49,997 exomes were available. Variants in *TTN* were annotated with the Ensembl Variant Effect Predictor version 95 with the --pick-allele flag^44^. LOFTEE was used to identify high-confidence loss of function variants in *TTN* (TTNtv): namely, stop-gained, splice-site disrupting, and frameshift variants^45^.

Of the samples with cardiac MRI that passed all of the disease exclusion and quality control measures described above, 11,878 also had exome sequencing. Of these, 53 samples had TTNtv in exons which are spliced into more than 90% of transcripts in the heart (“high-PSI”), which are likely to be relevant to cardiac phenotypes^5^.

We applied the the polygenic score most predictive of DCM (derived from the LVESVi) to the TTNtv carriers, and assessed whether that score was associated with LVESV, LVEDV, and LVEF in those individuals after adjustment for the first five principal components of ancestry, genotyping array, sex, and the cubic basis spline of age at enrollment. To minimize the risk of weak-instrument bias^46^, we re-computed weights for the best performing polygenic score (LVESVi) while excluding the 53 TTNtv carriers from the LVESVi GWAS. The original polygenic score and the recomputed score were similar (R^2^ = 0.9999 across the 53 samples with TTNtv). We defined statistical significance as a two-tailed P < (0.05 / 3 phenotypes) = 0.017.

## Acknowledgements

This work was conducted using the UK Biobank under application 7089.

## Data Availability

Data is made available from the UK Biobank to researchers from universities and other research institutions with genuine research inquiries following IRB and UK Biobank approval.

