## Supplement for "Analysis of cardiac magnetic resonance imaging traits in 29,000 individuals reveals shared genetic basis with dilated cardiomyopathy"

### Supplemental Material

#### Supplemental Tables

##### Supplemental Table 1: 96 curated disease phenotype definitions

| **Disease phenotype** | **Definition** |
|---|---|
| Hypertrophic cardiomyopathy | Self-reported history of hypertrophic cardiomyopathy during a verbal interview with a trained nurse; **or** hospitalization or death due to an ICD-10 code for hypertrophic cardiomyopathy (I42.1, I42.2); **or** hospitalization due to an ICD-9 code for hypertrophic cardiomyopathy (425.11, 425.18). |
| Heart failure | Self-reported history of heart failure or cardiomyopathy during verbal interview with trained nurse; **or** hospitalization for or death due to ICD-10 code for hypertensive heart disease, cardiomyopathy or heart failure (I11.0, I13.0, I13.2, I25.5, I42.0, I42.5, I42.8, I42.9, I50.0, I50.1, I50.9); **or** hospitalization due to ICD-9 code for heart failure or other primary cardiomyopathies (4254, 4280, 4281, 4289); **excluding** individuals with history of hypertrophic cardiomyopathy during verbal interview with trained nurse, or hospitalization for or death due to ICD-10 code for hypertrophic cardiomyopathy (I42.1, I42.2) |
| Nonischemic dilated cardiomyopathy | Hospitalization for or death due to ICD-10 code for dilated cardiomyopathy (I42.0); **excluding** individuals with history of coronary artery disease (as defined below), or history of hypertrophic cardiomyopathy during verbal interview with trained nurse, or hospitalization for or death due to ICD-10 code for hypertrophic cardiomyopathy (I42.1, I42.2) |
| Coronary artery disease | Self-reported history of myocardial infarction (MI), coronary artery bypass grafting, coronary artery angioplasty or triple heart bypass during verbal interview with trained nurse; **or** hospitalization for or death due to ICD-10 code for acute or subsequent myocardial infarction (I21, I22, I23, I24.1, I25.2); **or** hospitalization due to ICD-9 code for myocardial infarction (410, 411, 412); **or** hospitalization due to OPCS-4 code for coronary artery bypass grafting (K40, K41, K44, K45, K46), coronary endarterectomy (K47.1), or coronary angioplasty ± stenting (K49, K50.2, K75) |
| Stroke | Self-reported history of stroke during verbal interview with trained nurse; **or** hospitalization with or death due to ICD-10 code for nontraumatic subarachnoid hemorrhage, nontraumatic intracerebral |

| | |
|---|---|
| | hemorrhage, cerebral infarction, or unspecified stroke (I60-64); **or** hospitalization with or death due to ICD-9 code for subarachnoid hemorrhage, intracerebral hemorrhage, occlusion of cerebral arteries, or acute cerebrovascular disease (430, 431, 434, 436), as adjudicated centrally by the UK Biobank (http://biobank.ctsu.ox.ac.uk/crystal/refer.cgi?id=462) |
| Ischemic stroke | Self-reported history of ischemic stroke during verbal interview with trained nurse; **or** hospitalization with or death due to ICD-10 code for cerebral infarction, or unspecified stroke (I63, 64); **or** hospitalization with or death due to ICD-9 code for occlusion of cerebral arteries or acute cerebrovascular disease (434, 436), as adjudicated centrally by the UK Biobank (http://biobank.ctsu.ox.ac.uk/crystal/refer.cgi?id=462) |
| Intracerebral hemorrhage | Self-reported history of brain hemorrhage during verbal interview with trained nurse; **or** hospitalization with or death due to ICD-10 code for nontraumatic intracerebral hemorrhage (I61); **or** hospitalization with or death due to ICD-9 code for intracerebral hemorrhage (431), as adjudicated centrally by the UK Biobank (http://biobank.ctsu.ox.ac.uk/crystal/refer.cgi?id=462) |
| Subarachnoid hemorrhage | Self-reported history of subarachnoid hemorrhage during verbal interview with trained nurse; **or** hospitalization with or death due to ICD-10 code for nontraumatic subarachnoid hemorrhage (I60); **or** hospitalization with or death due to ICD-9 code for subarachnoid hemorrhage (430), as adjudicated centrally by the UK Biobank (http://biobank.ctsu.ox.ac.uk/crystal/refer.cgi?id=462) |
| Pulmonary hypertension | Hospitalization with or death due to ICD-10 code for primary or other secondary pulmonary hypertension (I27.0, I27.2); **or** hospitalization with ICD-9 code for primary pulmonary hypertension (4160) |
| Atrial fibrillation or flutter | Self-reported history of atrial fibrillation, atrial flutter, or cardioversion during verbal interview with trained nurse; **or** hospitalization with or death due to ICD-10 code for atrial fibrillation or atrial flutter (I48); **or** hospitalization with ICD-9 code for atrial fibrillation or atrial flutter (4273); **or** hospitalization with OPCS-4 code for percutaneous transluminal ablation (K57.1, K 62.1, K62.2, K62.3, K62.4) |
| Venous thromboembolism | Self-reported history of venous thromboembolic disease, pulmonary embolism or deep venous thrombosis during verbal interview with trained nurse; **or** hospitalization with or death due to ICD-10 code for pulmonary embolism (I26), phlebitis or thrombophlebitis (I80.0-I80.3, I80.8-I80.9), portal vein thrombosis (I81), Budd-Chiari syndrome (I82.0), or other coagulation defects (D68); **or** hospitalization with ICD-9 code for pulmonary embolism or phlebitis/thrombophlebitis (4151, 4511); **or** hospitalization with OPCS-4 code for insertion of inferior vena cava filter or open thrombectomy of lower extremity vein |

| | |
|---|---|
| | (L79.1, L90.2) |
| Peripheral artery disease | Self-reported history of peripheral vascular disease, arterial embolism, intermittent claudication, leg artery bypass, leg artery angioplasty, or leg amputation during verbal interview with trained nurse; **<u>or</u>** hospitalization with or death due to ICD-10 code for atherosclerosis of (non-coronary) arteries or peripheral vascular disease (I70.0, I70.00, I70.01, I70.2, I70.20, I70.21, I70.8, I70.80, I70.9, I70.90, I73.8 or I73.9); **<u>or</u>** hospitalization with ICD-9 code for atherosclerosis of arteries or peripheral vascular disease (4400, 4402, 4438, 4439); **<u>or</u>** hospitalization with OPCS-4 coded procedure for leg amputation, or leg artery procedure such as bypass, stent or angioplasty (X09.3-09.5, L21.6, L51.3, L51.6, L51.8, L52.1, L52.2, L54.1, L54.4, L54.8, L59.1-L59.8, L60.1, L60.2, L63.1, L63.5, L63.9, L66.7) |
| Hypertension | Self-reported history of hypertension, essential hypertension or high blood pressure during verbal interview with trained nurse; **<u>or</u>** hospitalization with or death due to ICD-10 code for essential hypertension, hypertensive heart disease, hypertensive renal disease, or secondary hypertension (I10, I11, I12, I13, I15); **<u>or</u>** hospitalization with ICD-9 code for essential hypertension, hypertensive heart disease, hypertensive renal disease, or secondary hypertension (401, 402, 403, 404, 405) |
| Hypercholesterolemia | Self-reported history of high cholesterol during verbal interview with trained nurse; **<u>or</u>** hospitalization with or death due to ICD-10 code for hypercholesterolemia, hypertriglyceridemia, or hyperlipidemia (E78.0-E78.2, E78.4, E78.5) |
| Supraventricular arrhythmia –general inclusive definition | Self-reported history of Wolff-Parkinson-White syndrome or supraventricular tachycardia during verbal interview with trained nurse; **<u>or</u>** hospitalization with or death due to ICD-10 code for Preexcitation syndrome, supraventricular tachycardia, atrial premature depolarization (I45.6, I47.1, I49.1); **<u>or</u>** hospitalization with ICD-9 code for anomalous atrioventricular excitation or paroxysmal supraventricular tachycardia (4267, 4270); **<u>or</u>** hospitalization with OPCS-4 coded procedure for open division of accessory pathway, percutaneous transluminal ablation of accessory pathway/atrial wall/conduction system (K52.4, K57.2, K57.4, K57.5) |
| Supraventricular arrhythmia –Wolff-Parkinson-White syndrome | Self-reported history of Wolff-Parkinson-White syndrome during verbal interview with trained nurse; **<u>or</u>** hospitalization with or death due to ICD-10 code for Preexcitation syndrome (I45.6); **<u>or</u>** hospitalization with ICD-9 for to anomalous atrioventricular excitation (4267); **<u>or</u>** hospitalization with OPCS-4 coded procedure for open division of accessory pathway, percutaneous transluminal ablation of accessory pathway (K52.4, K57.4) |

| | |
|---|---|
| Supraventricular arrhythmia –premature atrial contractions | Hospitalization with or death due to ICD-10 code for atrial premature depolarization (I49.1) |
| Supraventricular arrhythmia –supraventricular tachycardia | Self-reported history of supraventricular tachycardia during verbal interview with trained nurse; **or** hospitalization with or death due to ICD-10 code for supraventricular tachycardia (I47.1); **or** hospitalization with ICD-9 code for paroxysmal supraventricular tachycardia (4270); **or** hospitalization with OPCS-4 coded procedure for percutaneous transluminal ablation of atrial wall/conduction system (K57.2, K57.5) |
| Bradyarrhythmia – general inclusive definition | Self-reported history of sick sinus syndrome, pacemaker/defibrillator insertion, or pacemaker battery change during verbal interview with trained nurse; **or** hospitalization with or death due to ICD-10 code for atrioventricular and intraventricular block or sick sinus syndrome (I44, I45.0-I45.5, I49.5); **or** hospitalization with ICD-9 code for atrioventricular or intraventricular block (4260, 4261, 4263, 4264, 4265, 4266); **or** hospitalization with OPCS-4 coded procedure for cardiac pacemaker system (K60, K61) |
| Bradyarrhythmia – sinus node dysfunction | Self-reported history of sick sinus syndrome during verbal interview with trained nurse; **or** hospitalization with or death due to ICD-10 code for sick sinus syndrome (I49.5) |
| Bradyarrhythmia - AV block / distal conduction disease | Hospitalization with or death due to ICD-10 code for atrioventricular and intraventricular block (I44, I45.0-I45.5); **or** hospitalization with ICD-9 code for atrioventricular or intraventricular block (4260, 4261, 4263, 4264, 4265, 4266) |
| Bradyarrhythmia - Pacemaker | Self-reported history of pacemaker/defibrillator insertion, or pacemaker battery change during verbal interview with trained nurse; **or** hospitalization with OPCS-4 coded procedure for cardiac pacemaker system (K60, K61) |
| Implantable cardioverter defibrillator | Hospitalization with OPCS-4 coded procedure for cardioverter defibrillator introduced through the vein (K59) |
| Ventricular arrhythmia - General inclusive definition | Hospitalization with or death due to ICD-10 code for ventricular arrhythmias, ventricular premature depolarization or cardiac arrest (I46.0, I46.1, I46.9, I47.0, I47.2, I49.0, I49.3); **or** hospitalization with ICD-9 code for ventricular arrhythmias or cardiac arrest (4271, 4274, 4275); **or** hospitalization with OPCS-4 coded procedure for percutaneous transluminal/radiofrequency ablation (K57.6, K64.1) or resuscitation/defibrillation (X50.3, X50.4, X50.8, X50.9) |
| Ventricular | Hospitalization with or death due to ICD-10 code for ventricular |

| | |
|---|---|
| arrhythmia – Ventricular tachycardia | arrhythmias (I47.0, I47.2, I49.0); **or** hospitalization with ICD-9 code for ventricular arrhythmias (4271, 4274); **or** hospitalization with OPCS-4 coded procedure for percutaneous transluminal/radiofrequency ablation (K57.6, K64.1) |
| Ventricular arrhythmia – Ventricular premature depolarizations | Hospitalization with or death due to ICD-10 code for ventricular premature depolarization (I49.3) |
| Ventricular arrhythmia – Sudden cardiac death | Hospitalization with or death due to ICD-10 code for cardiac arrest (I46.0, I46.1, I46.9); **or** hospitalization with ICD-9 code for ventricular arrhythmias or cardiac arrest (4275); **or** hospitalization with OPCS-4 coded procedure for resuscitation/defibrillation (X50.3, X50.4, X50.8, X50.9) |
| Aortic valve disease | Self-reported history of aortic stenosis, aortic valve disease, aortic regurgitation or aortic valve repair/replacement during verbal interview with trained nurse; **or** hospitalization with or death due to ICD-10 code for rheumatic aortic valve disease (I06), unspecified aortic valve disorders (I08.0, I08.2, I08.3, I39.1), or nonrheumatic aortic valve disorders (I35); **or** hospitalization with ICD-9 code for rheumatic aortic insufficiency (3951), or unspecified diseases of aortic valve (3959, 4241); **or** hospitalization with OPCS-4 code for aortic valve repair/revision (K26, K30.2) |
| Aortic stenosis | Self-reported history of aortic stenosis during verbal interview with trained nurse; **or** hospitalization with or death due to ICD-10 code for rheumatic aortic stenosis (I06.0, I06.2) or nonrheumatic aortic stenosis (I35.0, I35.2) |
| Aortic regurgitation | Self-reported history of aortic regurgitation during verbal interview with trained nurse; **or** hospitalization with or death due to ICD-10 code for rheumatic aortic insufficiency (I06.1, I06.2) or nonrheumatic aortic insufficiency (I35.0, I35.2); **or** hospitalization with ICD-9 code for rheumatic aortic insufficiency (3951) |
| Mitral valve disease | Self-reported history of mitral valve disease, mitral stenosis, mitral valve prolapse, mitral regurgitation, or mitral valve repair/replacement during verbal interview with trained nurse; **or** hospitalization with or death due to ICD-10 code for rheumatic mitral valve diseases (I05), nonrheumatic mitral valve diseases (I34), or unspecified mitral valve disorders (I08.0, I08.1, I08.3, I39.0); **or** hospitalization with ICD-9 code for mitral stenosis ± insufficiency (3940, 3942), unspecified diseases of mitral valve (3949, 4240); **or** hospitalization with OPCS-4 code for mitral valve repair/revision (K25, K30.1), mitral valve annuloplasty (K34.1), percutaneous transluminal mitral valvotomy (K35.1) |

| | |
|---|---|
| Mitral stenosis | Self-reported history of mitral stenosis during verbal interview with trained nurse; **or** hospitalization with or death due to ICD-10 code for rheumatic and nonrheumatic mitral stenosis (I05.0, I34.2); **or** hospitalization with ICD-9 code for mitral stenosis ± insufficiency (3940, 3942); **or** hospitalization with OPCS-4 code for percutaneous transluminal mitral valvotomy (K35.1) |
| Mitral regurgitation | Self-reported history of mitral regurgitation during verbal interview with trained nurse; **or** hospitalization with or death due to ICD-10 code for rheumatic and nonrheumatic mitral insufficiency (I05.1, I05.2, I34.0); **or** hospitalization with ICD-9 code for mitral insufficiency (3942); **or** hospitalization with OPCS-4 code for mitral valve annuloplasty (K34.1) |
| Mitral valve prolapse | Self-reported history of mitral valve prolapse during verbal interview with trained nurse; **or** hospitalization with or death due to ICD-10 code for mitral valve prolapse (I34.1) |
| Tricuspid valve disease | Hospitalization with or death due to ICD-10 code for rheumatic tricuspid valve diseases (I07), nonrheumatic tricuspid valve diseases (I36), or unspecified tricuspid valve disorders (I08.1, I08.2, I08.3); **or** hospitalization with OPCS-4 code for tricuspid valve repair/revision (K27, K30.3), or tricuspid valve annuloplasty (K34.2) |
| Tricuspid stenosis | Hospitalization with or death due to ICD-10 code for rheumatic and nonrheumatic tricuspid stenosis (I07.0, I07.2, I36.0) |
| Tricuspid regurgitation | Hospitalization with or death due to ICD-10 code for rheumatic and nonrheumatic tricuspid regurgitation (I07.1, I07.2, I36.1); **or** hospitalization with OPCS-4 code for tricuspid valve annuloplasty (K34.2) |
| Pulmonary valve disease | Hospitalization with or death due to ICD-10 code for pulmonary valve disorders (I37, I39.3); **or** hospitalization with ICD-9 code for pulmonary valve disorders (4243); **or** hospitalization with OPCS-4 code for pulmonary valve repair/revision (K28, K30.4), or percutaneous transluminal pulmonary valve replacement (K35.7) |
| Pulmonary stenosis | Hospitalization with or death due to ICD-10 code for pulmonary stenosis (I37.0, I37.2) |
| Pulmonary regurgitation | Hospitalization with or death due to ICD-10 code for pulmonary insufficiency (I37.1, I37.2) |
| Valvular disease, unspecified | Self-reported history of heart valve problem/murmur/surgery or other valve repair/replacement during verbal interview with trained nurse; **or** hospitalization with or death due to ICD-10 code for multiple and/or unspecified rheumatic and nonrheumatic valve diseases (I08.8, I08.9, I09.1, I39.4); **or** hospitalization with OPCS-4 code for unspecified |

| | |
|---|---|
| | valve replacement/repair/procedure (K29, K34.3, K35.8) |
| Congenital heart disease | Hospitalization with or death due to ICD-10 code for congenital malformations of cardiac chambers/connexions, cardiac septa, aortic/mitral/tricuspid/pulmonary valves, and other/unspecified malformation (Q20.1, Q20.2, Q20.3, Q20.4, Q20.5, Q20.6, Q20.8, Q20.9; Q21, Q22, Q23, Q24); **or** hospitalization with ICD-9 code for congenital anomalies of the heart and cardiac septa (745, 746); **or** hospitalization with OPCS-4 code for repair of tetralogy of Fallot (K04), correction of total anomalous pulmonary venous connection (K07), repair of defect of atrioventricular/interatrial/interventricular/unspecified septum (K09, K10, K11, K12), transluminal repair of septal defect (K13), transluminal repair of atrial septum or patent oval foramen with prosthesis (K16.3, K16.5), creation of valved/other cardiac conduit (K18, K19, refashioning of atrium (K20) |
| Congenital heart disease – Ebstein's anomaly | Hospitalization with or death due to ICD-10 code for Ebstein's anomaly (Q22.5) |
| Diabetes mellitus, type 1 | Self-reported history of Type 1 diabetes during verbal interview with trained nurse; **or** hospitalization with or death due to ICD-10 code for insulin-dependent diabetes mellitus (E10) |
| Diabetes mellitus, type 2 | Self-reported history of type 2 diabetes during verbal interview with trained nurse; **or** hospitalization with or death due to ICD-10 code for non-insulin-dependent diabetes mellitus (E11) |
| Diabetes mellitus, all | Self-reported history of diabetes, gestational diabetes, type 1/type 2 or insulin use during verbal interview with trained nurse; **or** hospitalization with or death due to ICD-10 code for insulin-dependent and non-insulin-dependent diabetes mellitus (E10, E11), malnutrition-related diabetes mellitus (E12), other specified/unspecified diabetes mellitus (E13, E14); **or** hospitalization with ICD-9 code for diabetes mellitus with/without complications (2500, 2501, 2503, 2504, 2505, 2509) |
| Hyperthyroidism | Self-reported history of hyperthyroidism/thyrotoxicosis during verbal interview with trained nurse; **or** hospitalization with or death due to ICD-10 code for hyperthyroidism/thyrotoxicosis (E05); **or** hospitalization with ICD-9 code for thyrotoxicosis with/without goiter (242) |
| Hypothyroidism | Self-reported history of hypothyroidism/myxoedema during verbal interview with trained nurse; **or** hospitalization with or death due to ICD-10 code for hypothyroidism (E03); **or** hospitalization with ICD-9 code for acquired hypothyroidism (244) |

| | |
|---|---|
| Gout | Self-reported history of gout during verbal interview with trained nurse; **<u>or</u>** hospitalization with or death due to ICD-10 code for gout (M10.0, M10.2, M10.3, M10.4, M10.9); **<u>or</u>** hospitalization with ICD-9 code for gout (274) |
| Enlarged prostate | Self-reported history of enlarged prostate during verbal interview with trained nurse; **<u>or</u>** hospitalization with or death due to ICD-10 code for hyperplasia of prostate (N40); **<u>or</u>** hospitalization with ICD-9 code for hyperplasia of prostate (600) |
| Uterine fibroids | Self-reported history of uterine fibroids or myomectomy/fibroid removal during verbal interview with trained nurse; **<u>or</u>** hospitalization with or death due to ICD-10 code for leiomyoma of uterus (D25); **<u>or</u>** hospitalization with ICD-9 code for uterine leiomyoma (218) |
| Chronic kidney disease | Self-reported history of kidney failure ± dialysis, kidney nephropathy, IgA nephropathy, diabetic nephropathy or kidney transplant during verbal interview with trained nurse; **<u>or</u>** hospitalization with or death due to ICD-10 code for hypertensive renal disease, chronic renal failure, end stage renal failure or chronic kidney disease (I12.0, I13.1, I13.2, N18, N18.0-18.5, N18.8, N18.9); **<u>or</u>**<br>hospitalization with ICD-9 code due to chronic renal failure (585, 5859); **<u>or</u>**<br>hospitalization with OPCS-4 coded procedure for kidney transplantation (M01.1-01.5, M01.8, M01.9) |
| Gastroesophageal reflux disease | Self-reported history of gastroesophageal/gastric reflux during verbal interview with trained nurse; **<u>or</u>** hospitalization with or death due to ICD-10 code for gastroesophageal reflux disease (K21); **<u>or</u>** hospitalization with ICD-9 code for esophageal reflux (53010, 53011) |
| Irritable bowel syndrome | Self-reported history of irritable bowel syndrome during verbal interview with trained nurse; **<u>or</u>** hospitalization with or death due to ICD-10 code for irritable bowel syndrome (K58); **<u>or</u>** hospitalization with ICD-9 code for irritable colon (5641) |
| Cholelithiasis | Self-reported history of cholelithiasis/gallstones or gallstone removal during verbal interview with trained nurse; **<u>or</u>** hospitalization with or death due to ICD-10 code for gallstone ileus (K56.3) or cholelithiasis (K80; **<u>or</u>** hospitalization with ICD-9 code for cholelithiasis (574); **<u>or</u>** hospitalization with OPCS-4 coded procedure for open removal/percutaneous dissolution/fragmentation of gall bladder calculus (J21.1, J24.2, J24.3, J26.1) |
| Inflammatory bowel disease | Self-reported history of inflammatory bowel disease, Crohn’s disease, or ulcerative colitis during verbal interview with trained nurse; **<u>or</u>** hospitalization with or death due to ICD-10 code for Crohn’s disease or ulcerative colitis (K50, K51); **<u>or</u>** hospitalization with ICD-9 code for |

| | |
|---|---|
| | regional enteritis, idiopathic proctocolitis (555, 556) |
| Crohn’s disease | Self-reported history of Crohn’s disease during verbal interview with trained nurse; **or** hospitalization with or death due to ICD-10 code for Crohn’s disease (K50); **or** hospitalization with ICD-9 code for regional enteritis (555) |
| Ulcerative colitis | Self-reported history of ulcerative colitis during verbal interview with trained nurse; **or** hospitalization with or death due to ICD-10 code for ulcerative colitis (K51); **or** hospitalization with ICD-9 code for idiopathic proctocolitis (556) |
| Diverticular disease | Self-reported history of diverticular disease during verbal interview with trained nurse; **or** hospitalization with or death due to ICD-10 code for diverticular disease of intestine (K57); **or** hospitalization with ICD-9 code for diverticula of intestine (562) |
| Pancreatitis | Self-reported history of pancreatitis during verbal interview with trained nurse; **or** hospitalization with or death due to ICD-10 code for acute pancreatitis (K85) or alcohol-induced/other chronic pancreatitis (K86.0, K86.1); **or** hospitalization with ICD-9 code for acute/chronic pancreatitis (5770, 5771) |
| Migraine | Self-reported history of migraine during verbal interview with trained nurse; **or** hospitalization with or death due to ICD-10 code for migraine (G43) |
| Depression | Self-reported history of depression during verbal interview with trained nurse; **or** hospitalization with or death due to ICD-10 code for depressive episode or recurrent depressive disorder (F32, F33); **or** hospitalization with ICD-9 code for depressive disorder (3119) |
| Bipolar disorder | Self-reported history of mania/bipolar disorder/manic depression during verbal interview with trained nurse; **or** hospitalization with or death due to ICD-10 code for bipolar affective disorder (F31) |
| Anxiety | Self-reported history of anxiety/panic attacks during verbal interview with trained nurse; **or** hospitalization with or death due to ICD-10 code for anxiety disorders (F41) |
| Schizophrenia | Self-reported history of schizophrenia during verbal interview with trained nurse; **or** hospitalization with or death due to ICD-10 code for schizophrenia (F20) |
| Post-traumatic stress disorder | Self-reported history of post-traumatic stress disorder during verbal interview with trained nurse; **or** hospitalization with or death due to ICD-10 code for post-traumatic stress disorder (F43.1) |

| | |
|---|---|
| Multiple sclerosis | Self-reported history of multiple sclerosis during verbal interview with trained nurse; **<u>or</u>** hospitalization with or death due to ICD-10 code for multiple sclerosis (G35); **<u>or</u>** hospitalization with ICD-9 code for multiple sclerosis (3409) |
| Parkinson's disease | Self-reported history of Parkinson's disease during verbal interview with trained nurse; **<u>or</u>** hospitalization with or death due to ICD-10 code for Parkinson's disease (G20) or dementia in Parkinson's disease (F02.3); **<u>or</u>** hospitalization with ICD-9 code for paralysis agitans (3320) |
| Epilepsy | Self-reported history of epilepsy during verbal interview with trained nurse; **<u>or</u>** hospitalization with or death due to ICD-10 code for epilepsy (G40); **<u>or</u>** hospitalization with ICD-9 code for epilepsy (3450, 3451, 3452, 3454, 3459) |
| Alzheimer's / Dementia | Self-reported history of dementia/Alzheimer's/cognitive impairment during verbal interview with trained nurse; **<u>or</u>** hospitalization with or death due to ICD-10 code for dementia in Alzheimer's disease (F00) |
| Back pain | Self-reported history of back pain or sciatica during verbal interview with trained nurse; **<u>or</u>** hospitalization with or death due to ICD-10 code for Dorsalgia (M54); **<u>or</u>** hospitalization with ICD-9 code for cervicalgia, spinal stenosis, pain in thoracic spine, lumbago, sciatica, thoracic or lumbosacral neuritis/radiculitis, unspecified backache (7231, 7240, 7241, 7242, 7243, 7244, 7245) |
| Sciatica | Self-reported history of sciatica during verbal interview with trained nurse; **<u>or</u>** hospitalization with or death due to ICD-10 code for sciatica (M54.3); **<u>or</u>** hospitalization with ICD-9 code for sciatica (7243) |
| Osteoporosis | Self-reported history of osteoporosis during verbal interview with trained nurse; **<u>or</u>** hospitalization with or death due to ICD-10 code for osteoporosis with/without pathological fracture (M80, M81); **<u>or</u>** hospitalization with ICD-9 code for osteoporosis (7330) |
| Intervertebral disc displacement | Self-reported history of prolapsed/slipped disc during verbal interview with trained nurse; **<u>or</u>** hospitalization with or death due to ICD-10 code for other cervical/intervertebral disc displacement (M50.2, M51.2); **<u>or</u>** hospitalization with ICD-9 code for cervical/thoracic/lumbar/unspecified intervertebral disc displacement (7220, 7221, 7222) |
| Osteoarthritis | Self-reported history of osteoarthritis during verbal interview with trained nurse; **<u>or</u>** hospitalization with or death due to ICD-10 code for polyarthrosis, coxarthrosis, gonarthrosis, first carpometacarpal or other arthrosis (M15, M16, M17, M18, M19); **<u>or</u>** hospitalization with ICD-9 code for osteoarthrosis (715) |
| Rheumatoid arthritis | Self-reported history of rheumatoid arthritis during verbal interview with |

| | |
|---|---|
| | trained nurse; **or** hospitalization with or death due to ICD-10 code for rheumatoid arthritis (M05, M06); **or** hospitalization with ICD-9 code for rheumatoid arthritis (714) |
| Lupus erythematosus | Self-reported history of systemic lupus erythematosus during verbal interview with trained nurse; **or** hospitalization with or death due to ICD-10 code for lupus erythematosus (L93) or systemic lupus erythematosus (M32.1, M32.8, M32.9); **or** hospitalization with ICD-9 code for lupus erythematosus (6954) |
| Sarcoidosis | Self-reported history of sarcoidosis during verbal interview with trained nurse; **or** hospitalization with or death due to ICD-10 code for sarcoidosis (D86); **or** hospitalization with ICD-9 code for sarcoidosis (135) |
| Psoriasis | Self-reported history of psoriasis during verbal interview with trained nurse; **or** hospitalization with or death due to ICD-10 code for psoriasis (L40), psoriatic arthropathy, arthritis mutilans, psoriatic spondylitis (M07.0, M07.1, M07.2, M07.3); **or** hospitalization with ICD-9 code for psoriasis/psoriatic arthropathy (6960, 6961) |
| Dermatitis | Self-reported history of eczema/dermatitis during verbal interview with trained nurse; **or** hospitalization with or death due to ICD-10 code for atopic dermatitis, seborrhoeic dermatitis, diaper dermatitis, allergic contact dermatitis, irritant/unspecified contact dermatitis (L20, L21, L22, L23, L24, L25, L26, L27, L30), Lichen simplex chronicus and prurigo (L28), pruritus (L29); **or** hospitalization with ICD-9 code for atopic/contact dermatitis (691, 692) |
| Iron deficiency anemia | Self-reported history of iron deficiency anemia during verbal interview with trained nurse; **or** hospitalization with or death due to ICD-10 code for iron deficiency anemia (D50); **or** hospitalization with ICD-9 code for iron deficiency anemia (280) |
| Asthma | Self-reported history of asthma during verbal interview with trained nurse; **or** hospitalization with or death due to ICD-10 code for asthma or status asthmaticus (J45, J46); **or** hospitalization with ICD-9 code for asthma (493) |
| Chronic obstructive pulmonary disease | Self-reported history of chronic obstructive airways/emphysema during verbal interview with trained nurse; **or** hospitalization with or death due to ICD-10 code for chronic bronchitis, emphysema, or other chronic obstructive pulmonary disease (J41, J42, J43, J44); **or** hospitalization with ICD-9 code for chronic bronchitis, emphysema or unspecified chronic airways obstruction (491, 492, 496) |
| Pneumonia | Self-reported history of pneumonia during verbal interview with trained nurse; **or** hospitalization with or death due to ICD-10 code for |

| | |
|---|---|
| | pneumonia (J12, J13, J14, J15, J16, J17, J18); **or** hospitalization with ICD-9 code for pneumonia (481, 482, 483, 484, 485, 486) |
| Allergic rhinitis | Self-reported history of hayfever/allergic rhinitis during verbal interview with trained nurse; **or** hospitalization with or death due to ICD-10 code for allergic rhinitis (J30.1-J30.4); **or** hospitalization with ICD-9 code for allergic rhinitis (477) |
| Sleep apnea | Self-reported history of sleep apnea during verbal interview with trained nurse; **or** hospitalization with or death due to ICD-10 code for sleep apnea (G47.3) |
| Cataract | Self-reported history of cataract during verbal interview with trained nurse; **or** hospitalization with or death due to ICD-10 code for cataract (H25, H26); **or** hospitalization with ICD-9 code for cataract (366) |
| Glaucoma | Self-reported history of glaucoma during verbal interview with trained nurse; **or** hospitalization with or death due to ICD-10 code for glaucoma (H40); **or** hospitalization with ICD-9 code for glaucoma (365) |
| Lung cancer | Self-reported history of lung cancer during verbal interview with trained nurse; **or** hospitalization with or death due to ICD-10 code for malignant neoplasm of bronchus and lung (C34); **or** hospitalization with ICD-9 code for malignant neoplasm of bronchus and lung (1629) |
| Breast cancer | Self-reported history of breast cancer during verbal interview with trained nurse; **or** hospitalization with or death due to ICD-10 code for malignant neoplasm of breast (C50); **or** hospitalization with ICD-9 code for malignant neoplasm of breast (174) |
| Colorectal cancer | Self-reported history of colorectal/sigmoid/rectal cancer during verbal interview with trained nurse; **or** hospitalization with or death due to ICD-10 code for malignant neoplasm of colon (C18.0, C18.2-18.9); **or** hospitalization with ICD-9 code for malignant neoplasm of colon/rectum (1532, 1533, 1541) |
| Skin cancer | Self-reported history of skin, malignant melanoma, non-melanoma skin cancer, or squamous cell carcinoma during verbal interview with trained nurse; **or** hospitalization with or death due to ICD-10 code for malignant melanoma or other malignant neoplasms of skin (C43, C44); **or** hospitalization with ICD-9 code for malignant melanoma/neoplasm of skin (172, 173) |
| Prostate cancer | Self-reported history of prostate cancer during verbal interview with trained nurse; **or** hospitalization with or death due to ICD-10 code for malignant neoplasm of prostate (C61) |
| Cervical cancer | Self-reported history of cervical cancer during verbal interview with |

| | |
|---|---|
| | trained nurse; **or** hospitalization with or death due to ICD-10 code for malignant neoplasm of cervix uteri (C53) |
| Bladder cancer | Self-reported history of bladder cancer during verbal interview with trained nurse; **or** hospitalization with or death due to ICD-10 code for malignant neoplasm of bladder (C67); **or** hospitalization with ICD-9 code for malignant neoplasm of bladder (188) |
| Gestational hypertension – preeclampsia | Self-reported history of gestational hypertension/pre-eclampsia during verbal interview with trained nurse; **or** hospitalization with or death due to ICD-10 code for pre-existing hypertensive disorder with superimposed proteinuria (O11), gestational hypertension (O13, O14), eclampsia (O15); **or** hospitalization with ICD-9 code for transient hypertension of pregnancy, preeclampsia or eclampsia (6423, 6424, 6425, 6426, 6427) |

#### Supplemental Table 2: Clinical characteristics of UK Biobank participants with cardiac MRI data

| | Mean or N (%) | SD | Mean or N (%) | SD | Mean or N (%) | SD |
|---|---|---|---|---|---|---|
| | All participants | | Women | | Men | |
| **Participants** | 29041 | | 15282 (52.6%) | | 13759 (47.4%) | |
| **Ancestry** | | | | | | |
| **African** | 119 (0.4%) | | 58 (0.4%) | | 61 (0.4%) | |
| **East Asian** | 87 (0.3%) | | 52 (0.3%) | | 35 (0.3%) | |
| **European** | 28189 (97.1%) | | 14868 (97.3%) | | 13327 (96.9%) | |
| **South Asian** | 243 (0.8%) | | 83 (0.5%) | | 160 (1.2%) | |
| **Other** | 397 (1.3%) | | 221 (1.4%) | | 176 (1.3%) | |
| **Age (Years)** | 63.6 | 7.5 | 63.1 | 7.4 | 64.2 | 7.6 |
| **BMI ($kg/m^2$)** | 26.5 | 4.2 | 26.1 | 4.5 | 27 | 3.7 |
| **Height (cm)** | 169.4 | 9.2 | 163.1 | 6.1 | 176.4 | 6.5 |
| **Weight (kg)** | 76.3 | 14.6 | 69.2 | 12.4 | 84.2 | 12.8 |
| **Systolic blood pressure (mmHg)** | 135.7 | 16.6 | 132.3 | 17.1 | 139.4 | 15.2 |
| **Diastolic blood pressure (mmHg)** | 80.7 | 9.2 | 78.8 | 9.1 | 82.8 | 8.9 |
| **Self-reported activity level (MET-min/week)** | 2660 | 2954 | 2567 | 2818 | 2763 | 3096 |
| **Wore actigraph** | 13223 (45.5%) | | 7238 (47.4%) | | 5985 (43.5%) | |
| **Average acceleration (milli-gravities)** | 28.2 | 9.2 | 28.6 | 8.7 | 27.8 | 9.8 |

| | | | | | | |
|---|---|---|---|---|---|---|
| **Alcohol consumption (standard drinks/week)** | 8.9 | 9.1 | 6.4 | 6.7 | 11.7 | 10.4 |
| **Currently using tobacco** | 1813 (6.2%) | | 780 (5.1%) | | 1033 (7.5%) | |
| **Ever used tobacco** | 7149 (24.6%) | | 3425 (22.4%) | | 3724 (27.1%) | |
| **Pack years** | 4.6 | 11 | 3.6 | 9 | 5.6 | 12.7 |
| **Type 2 diabetes** | 719 (2.5%) | | 266 (1.7%) | | 453 (3.3%) | |
| **Hypertension** | 7291 (25.1%) | | 3185 (20.8%) | | 4106 (29.8%) | |
| **Hyperlipidemia** | 4569 (15.7%) | | 1785 (11.7%) | | 2784 (20.2%) | |
| **LVEDV (mL)** | 136.9 | 27.7 | 122.6 | 19.5 | 152.8 | 26.8 |
| **LVEDVi (mL/m$^2$)** | 72.4 | 11.8 | 69.6 | 10.2 | 75.6 | 12.7 |
| **LVESV (mL)** | 48.8 | 15.2 | 41.1 | 10.3 | 57.4 | 15.2 |
| **LVESVi (mL/m$^2$)** | 25.7 | 6.9 | 23.3 | 5.6 | 28.3 | 7.2 |
| **LVEF (%)** | 64.9 | 5.5 | 66.7 | 4.8 | 62.8 | 5.5 |
| **SV (mL)** | 88.1 | 15.8 | 81.5 | 12.2 | 95.4 | 16.1 |
| **SVi (mL/m$^2$)** | 46.7 | 7.1 | 46.3 | 6.5 | 47.2 | 7.8 |

### Supplemental Table 3: LD score regression

| | **Lambda** | **Intercept** |
|---|---|---|
| **LVEDV** | 1.146 | 1.026 |
| **LVEDVi** | 1.096 | 1.002 |
| **LVESV** | 1.253 | 1.103 |
| **LVESVi** | 1.096 | 0.987 |
| **LVEF** | 1.096 | 1.002 |
| **SV** | 1.096 | 1.02 |
| **SVi** | 1.096 | 1.008 |

Lambda represents the genomic inflation factor. Intercept represents the intercept from LD score regression.

### Supplemental Table 4: Genes in cardiomyopathy panels

The 129 genes in this table were assembled from 3 commercially available cardiomyopathy gene panels noted in the main text: from GeneDx, Invitae, and the Partners Laboratory for Molecular Medicine.

| | | | | | | | | | |
|---|---|---|---|---|---|---|---|---|---|
| A2ML1 | CALR3 | DOLK | GAA | LAMP2 | MT-TH | MYH7 | PDLIM3 | SCN5A | TMEM70 |
| ABCC9 | CASQ2 | DSC2 | GATA4 | LDB3 | MT-TI | MYL2 | PKP2 | SDHA | TMPO |
| ACADVL | CAV3 | DSG2 | GATA6 | LMNA | MT-TK | MYL3 | PLEKHM2 | SGCD | TNNC1 |
| ACTC1 | CBL | DSP | GATAD | LRRC10 | MT-TL1 | MYLK2 | PLN | SHOC2 | TNNI3 |

| | | | | | | | | | |
|---|---|---|---|---|---|---|---|---|---|
| | | | 1 | | | | | | |
| ACTN2 | CHRM2 | DTNA | GLA | MAP2K1 | MT-TL2 | MYOM1 | PRDM16 | SLC22A5 | TNNT2 |
| AGL | CPT2 | ELAC2 | HCN4 | MAP2K2 | MT-TM | MYOZ2 | PRKAG2 | SOS1 | TOR1AIP1 |
| AKAP9 | CRYAB | EMD | HFE | MED12 | MT-TQ | MYPN | PTPN11 | SOS2 | TPM1 |
| ALMS1 | CSRP3 | EYA4 | HRAS | MIB1 | MT-TS1 | NEBL | RAF1 | SPRED1 | TRDN |
| ALPK3 | CTF1 | FHL1 | ILK | MT-ND1 | MT-TS2 | NEXN | RASA1 | TAZ | TTN |
| ANKRD1 | CTNNA3 | FHL2 | JPH2 | MT-ND5 | MTO1 | NF1 | RBM20 | TBX20 | TTR |
| BAG3 | DES | FKRP | JUP | MT-ND6 | MURC | NKX2-5 | RIT1 | TCAP | TXNRD2 |
| BRAF | DMD | FKTN | KRAS | MT-TD | MYBPC3 | NPPA | RRAS | TGFB3 | VCL |
| CACNA1C | DNAJC19 | FLNC | LAMA4 | MT-TG | MYH6 | NRAS | RYR2 | TMEM43 | |

### Supplemental Table 5: Colocation with Mendelian cardiomyopathy genes

| | | | | |
|---|---|---|---|---|
| ACTN2 | FLNC | MYL2 | PTPN11 | TTN |
| ALPK3 | KRAS | PLEKHM2 | RYR2 | |
| BAG3 | MYH6 | PLN | TMEM43 | |
| FKRP | MYH7 | PRDM16 | TNNC1 | |

Each listed gene is located within 1 megabase of a lead SNP from one of the seven traits, and was found in the list of 129 Mendelian cardiomyopathy-related genes in **Supplemental Table 3**.

### Supplemental Table 6: PheWAS results for curated phenotypes

| **Curated Phenotype** | **PRS** | **N_People** | **N With Phenotype** | **Beta** | **SE** | **P** |
|---|---|---|---|---|---|---|
| Hypothyroidism | sv | 456237 | 29047 | -7.05E-02 | 6.14E-03 | 1.38E-30 |
| Hypertension | sv | 456237 | 158721 | -3.43E-02 | 3.26E-03 | 6.52E-26 |
| Dilated cardiomyopathy | lvesvi | 456237 | 708 | 3.86E-01 | 3.85E-02 | 1.32E-23 |
| Dilated cardiomyopathy | lvesv | 456237 | 708 | 3.57E-01 | 3.79E-02 | 4.73E-21 |
| Dilated cardiomyopathy | lvef | 456237 | 708 | -3.47E-01 | 3.99E-02 | 3.54E-18 |
| Heart Failure | lvef | 456237 | 9245 | -9.50E-02 | 1.09E-02 | 3.97E-18 |
| Heart Failure | lvesv | 456237 | 9245 | 8.89E-02 | 1.06E-02 | 5.72E-17 |
| Heart Failure | lvesvi | 456237 | 9245 | 8.91E-02 | 1.07E-02 | 9.33E-17 |
| Hypothyroidism | lvedvi | 456237 | 29047 | -4.98E-02 | 6.14E-03 | 4.90E-16 |
| Hypertension | svi | 456237 | 158721 | -2.62E-02 | 3.27E-03 | 1.20E-15 |
| Dilated cardiomyopathy | lvedv | 456237 | 708 | 2.91E-01 | 3.86E-02 | 5.50E-14 |
| Hypothyroidism | svi | 456237 | 29047 | -4.76E-02 | 6.19E-03 | 1.50E-14 |
| Hypertrophic cardiomyopathy | lvef | 456237 | 418 | 3.08E-01 | 4.91E-02 | 3.50E-10 |
| Dilated cardiomyopathy | lvedvi | 456237 | 708 | 2.25E-01 | 3.83E-02 | 3.95E-09 |
| Hypothyroidism | lvedv | 456237 | 29047 | -3.56E-02 | 6.16E-03 | 7.25E-09 |

| | | | | | | |
|---|---|---|---|---|---|---|
| Hypertension | lvef | 456237 | 158721 | -1.83E-02 | 3.34E-03 | 4.00E-08 |
| Coronary Artery Disease | sv | 456237 | 25396 | -3.58E-02 | 6.62E-03 | 6.42E-08 |
| Hypertrophic cardiomyopathy | lvesv | 456237 | 418 | -2.63E-01 | 4.90E-02 | 7.94E-08 |
| Hypertrophic cardiomyopathy | lvesvi | 456237 | 418 | -2.71E-01 | 4.92E-02 | 3.74E-08 |
| Hypertension | lvedvi | 456237 | 158721 | -1.62E-02 | 3.25E-03 | 6.19E-07 |
| Mitral regurgitation | lvesvi | 456237 | 3405 | 8.77E-02 | 1.74E-02 | 4.92E-07 |
| Mitral valve disease | lvef | 456237 | 5995 | -7.11E-02 | 1.35E-02 | 1.26E-07 |
| Myocardial Infarction | lvef | 456237 | 19437 | -3.75E-02 | 7.67E-03 | 9.97E-07 |
| Myocardial Infarction | sv | 456237 | 19437 | -3.71E-02 | 7.46E-03 | 6.56E-07 |
| Coronary Artery Disease | lvef | 456237 | 25396 | -3.56E-02 | 6.81E-03 | 1.74E-07 |
| Atrial fibrillation or flutter | lvedvi | 456237 | 21493 | -3.50E-02 | 7.10E-03 | 8.63E-07 |
| Atrial fibrillation or flutter | lvesvi | 456237 | 21493 | -3.68E-02 | 7.17E-03 | 2.81E-07 |
| Implantable cardioverter defibrillator | lvesv | 456237 | 1051 | 1.54E-01 | 3.10E-02 | 6.76E-07 |
| Heart Failure | lvedv | 456237 | 9245 | 5.03E-02 | 1.07E-02 | 2.46E-06 |
| Mitral regurgitation | lvef | 456237 | 3405 | -8.31E-02 | 1.78E-02 | 3.06E-06 |
| Mitral regurgitation | lvesv | 456237 | 3405 | 7.84E-02 | 1.73E-02 | 5.77E-06 |
| Mitral valve disease | lvesvi | 456237 | 5995 | 6.24E-02 | 1.32E-02 | 2.19E-06 |
| Coronary Artery Disease | lvedvi | 456237 | 25396 | -3.09E-02 | 6.63E-03 | 3.01E-06 |
| Coronary Artery Disease | svi | 456237 | 25396 | -3.04E-02 | 6.65E-03 | 4.97E-06 |
| Supraventricular arrhythmia General inclusive definition clinical | svi | 456237 | 4080 | -7.02E-02 | 1.58E-02 | 8.72E-06 |
| Psoriasis | sv | 456237 | 7096 | -4.82E-02 | 1.19E-02 | 5.10E-05 |
| Hypertension | lvesv | 456237 | 158721 | 1.35E-02 | 3.26E-03 | 3.62E-05 |
| Mitral valve disease | lvesv | 456237 | 5995 | 5.22E-02 | 1.31E-02 | 6.51E-05 |
| Myocardial Infarction | svi | 456237 | 19437 | -3.21E-02 | 7.49E-03 | 1.80E-05 |
| Tricuspid valve disease | lvef | 456237 | 2624 | -7.96E-02 | 2.03E-02 | 8.52E-05 |
| ischemic cardiomyopathy | lvesv | 456237 | 411 | 1.94E-01 | 4.95E-02 | 8.80E-05 |
| Atrial fibrillation or flutter | lvesv | 456237 | 21493 | -2.90E-02 | 7.12E-03 | 4.73E-05 |
| Atrial fibrillation or flutter | sv | 456237 | 21493 | -2.98E-02 | 7.09E-03 | 2.68E-05 |
| Supraventricular arrhythmia General inclusive definition clinical | sv | 456237 | 4080 | -6.32E-02 | 1.57E-02 | 5.36E-05 |

Effect size, standard error, and P value are displayed for the association between manually curated phenotypes and the seven cardiac trait polygenic scores. Associations with $P < 1 \times 10^{-5}$ with any of the 7 polygenic scores are shown.

### Supplemental Table 7: Characteristics of unrelated participants who did not undergo cardiac MRI

| | **Mean or N (%)** | **SD** | **Unit** |
|---|---|---|---|
| **Participants** | 358556 | | |

| | | | |
|---|---|---|---|
| **Male** | 157621 (44.0%) | | |
| **Ancestry** | | | |
| **African** | 2923 (0.8%) | | |
| **East Asian** | 1307 (0.4%) | | |
| **European** | 344708 (96.1%) | | |
| **South Asian** | 5994 (1.7%) | | |
| **Other** | 3624 (1.0%) | | |
| **Birth Year** | 1952 | 8 | **Years** |
| **BMI** | 27.4 | 4.8 | **kg/m^2** |
| **Height** | 168.3 | 9.3 | **cm** |
| **Weight** | 77.7 | 15.9 | **kg** |
| **Systolic blood pressure** | 138 | 18.6 | **mmHg** |
| **Diastolic blood pressure** | 82.4 | 10.1 | **mmHg** |
| **Alcohol consumption** | 8.4 | 10.7 | **Standard drinks/week** |
| **Activity level** | 2675 | 3777 | **MET-min/week** |
| **Currently using tobacco** | 38501 (10.7%) | | |
| **Type 2 diabetes** | 15357 (4.3%) | | |
| **Hypertension** | 95773 (26.7%) | | |
| **Hyperlipidemia** | 53369 (14.9%) | | |

This table presents the characteristics of the unrelated participants with genetic data who did not undergo cardiac MRI, and who were assessed for the ability of the 19-SNP score to predict incident DCM.

### Supplemental Table 8: Polygenic scores from cardiac MRI phenotypes are associated with incident DCM

| Source of SNP Score | Hazard Ratio | 95% CI Lower | 95% CI Upper | P-value | SNPs |
|---|---|---|---|---|---|
| LVEDV | 1.37 | 1.24 | 1.52 | 1.31E-09 | 14 |
| LVESV | 1.49 | 1.35 | 1.65 | 8.80E-15 | 24 |
| LVEF | 0.69 | 0.62 | 0.76 | 3.41E-12 | 16 |
| SV | 1.1 | 0.99 | 1.21 | 6.93E-02 | 3 |
| LVEDVi | 1.26 | 1.14 | 1.40 | 7.48E-06 | 8 |
| **LVESVi** | **1.54** | **1.39** | **1.70** | **2.05E-16** | **19** |
| SVi | 1.02 | 0.92 | 1.13 | 6.85E-01 | 6 |

"Hazard Ratio" represents the hazard ratio of a one standard deviation increase in the SNP score on the probability of developing dilated cardiomyopathy.

### Supplemental Table 9: Characteristics of TTNtv carriers

| | Mean or N (%) | SD | Unit |
|---|---|---|---|
| **Participants** | 53 | | |
| **Male** | 20 (37.7%) | | |
| **Ancestry** | | | |
| **European** | 53 (100%) | | |
| **Non-European** | 0 (0%) | | |
| **Age at Enrollment** | 54.7 | 7.9 | years |
| **BMI** | 27.1 | 4.2 | kg/m^2 |
| **Height** | 168 | 9.9 | cm |
| **Weight** | 76.2 | 12.6 | kg |

### Supplemental Figures

#### Supplemental Figure 1: Flow diagram

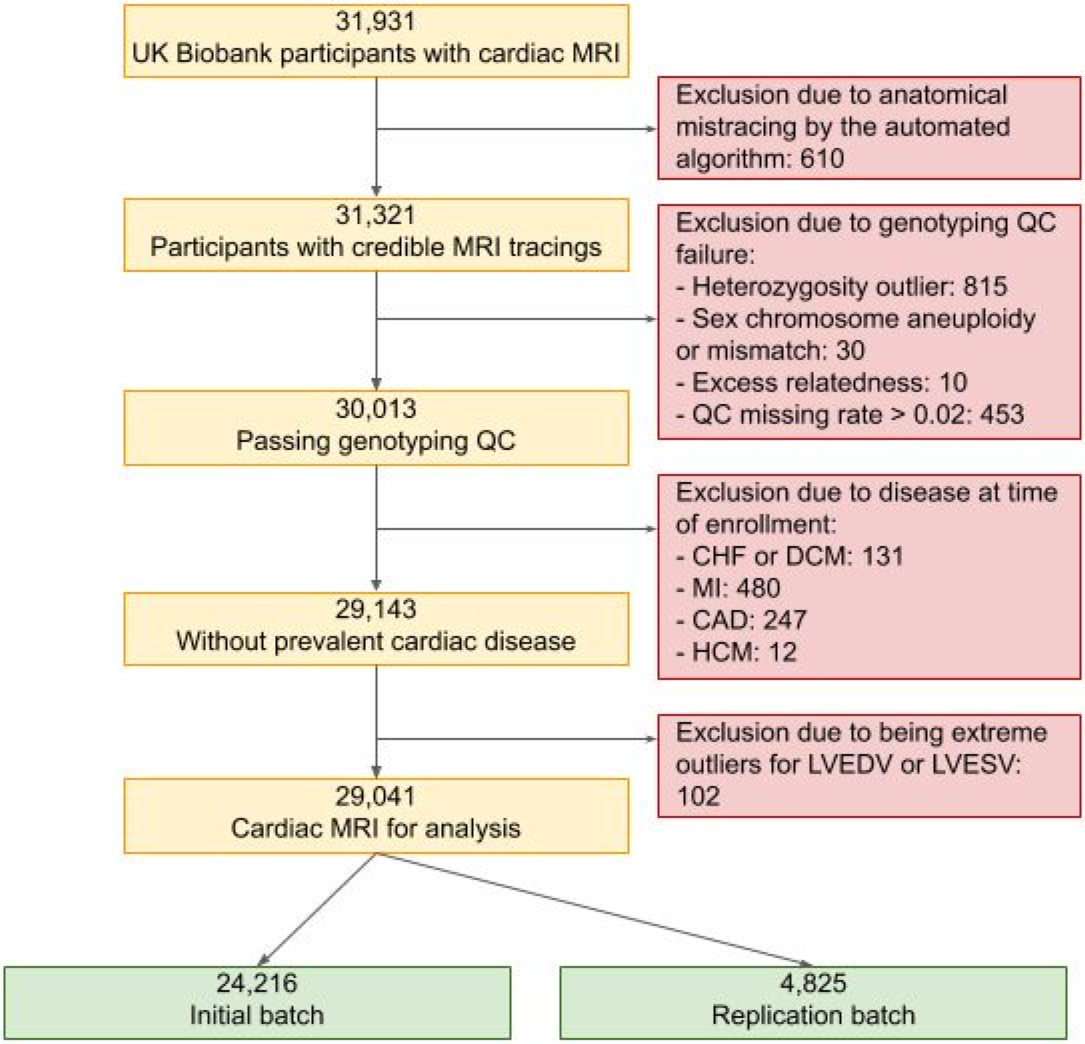

Supplemental Figure 2: Sex-stratified distributions of cardiac MRI phenotypes

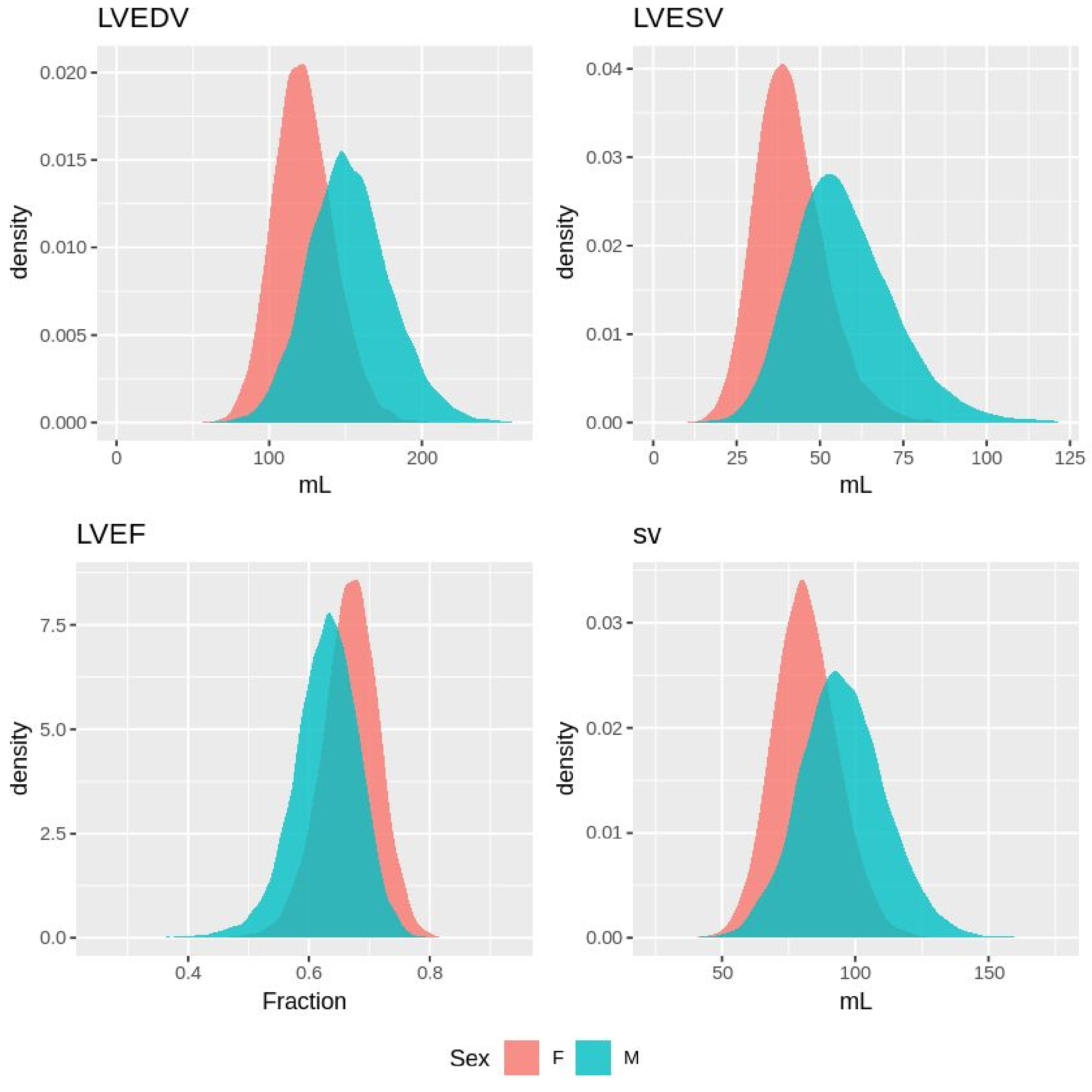


The population distribution of each of the non-BSA-indexed cardiac MRI phenotypes is represented through a density plot. Men and women are displayed separately, because the mean and variance for each phenotype vary by sex.

#### Supplemental Figure 3: Cross-trait correlation

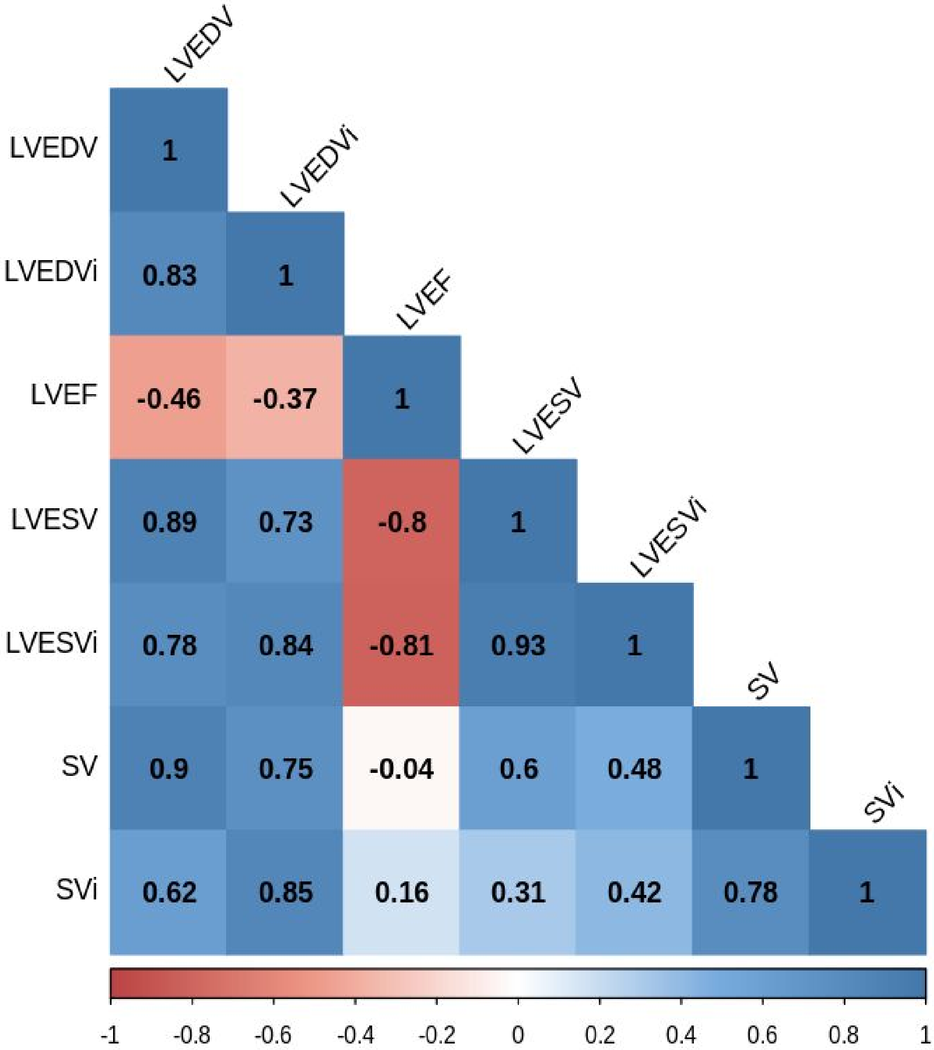


For each cardiac MRI phenotype pair, the Pearson correlation is represented by a number ranging from -1 (perfect anticorrelation) to 1 (perfect correlation), representing the degree to which each trait is correlated with another.

#### Supplemental Figure 4: Genetic correlation

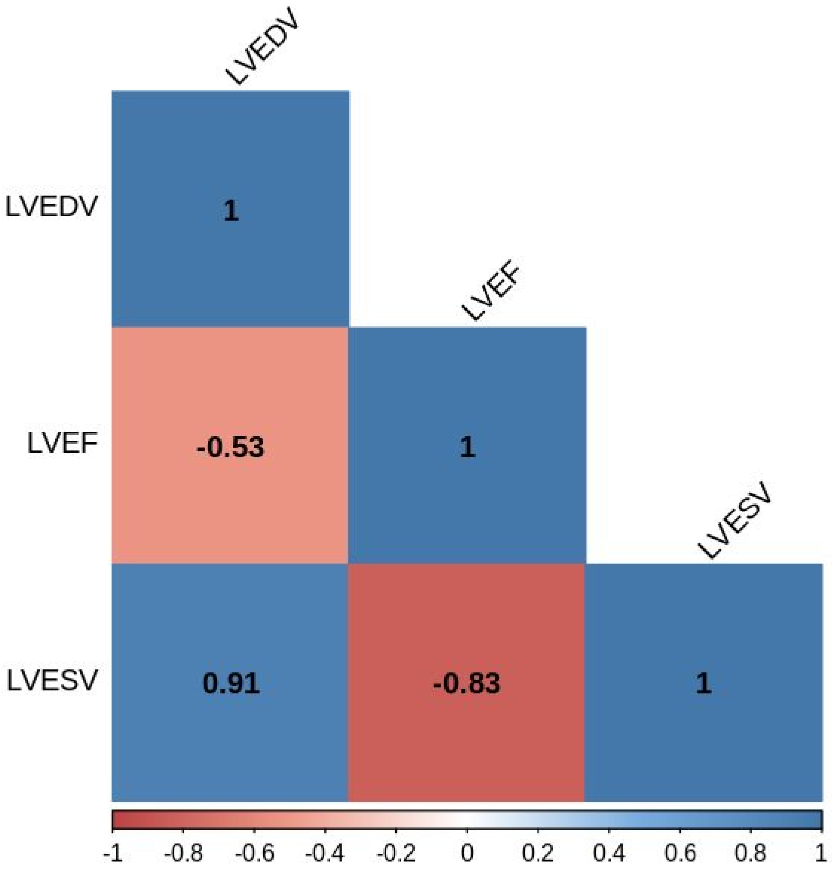


For each cardiac MRI phenotype pair, the genetic correlation as measured by BOLT REML is represented by a number ranging from -1 (perfect anticorrelation) to 1 (perfect correlation).

#### Supplemental Figure 5: MAGMA gene expression enrichment

For each cardiac phenotype, the associations of gene expression sets from GTEx were evaluated using MAGMA, running on the FUMA platform v1.3.4b. Gene expression sets significantly enriched after Bonferroni correction are highlighted in red.

#### LVEDV

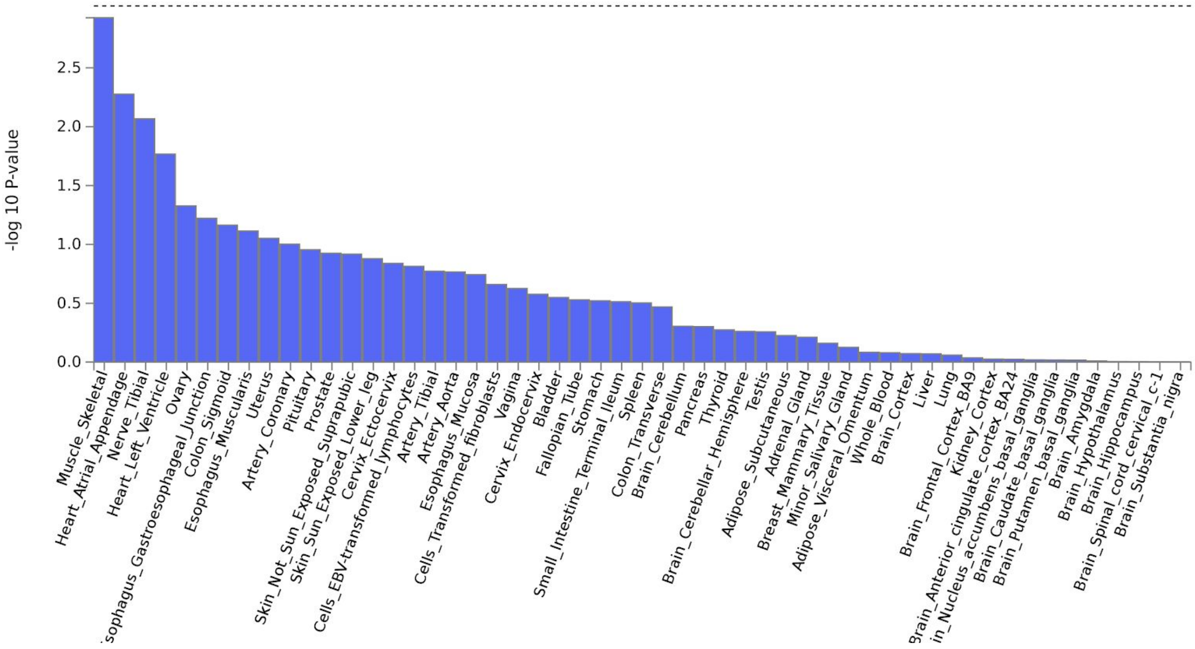
-log 10 P-value
0.0
0.5
1.0
1.5
2.0
2.5
Muscle_Skeletal
Heart_Atrial_Appendage
Nerve_Tibial
Heart_Left_Ventricle
Ovary
sophagus_Gastroesophageal_Junction
Colon_Sigmoid
Esophagus_Muscularis
Uterus
Artery_Coronary
Pituitary
Prostate
Skin_Not_Sun_Exposed_Suprapubic
Skin_Sun_Exposed_Lower_leg
Cervix_Ectocervix
Cells_EBV-transformed_lymphocytes
Artery_Tibial
Artery_Aorta
Esophagus_Mucosa
Cells_Transformed_fibroblasts
Vagina
Cervix_Endocervix
Bladder
Fallopian_Tube
Stomach
Small_Intestine_Terminal_Ileum
Spleen
Colon_Transverse
Brain_Cerebellum
Pancreas
Thyroid
Brain_Cerebellar_Hemisphere
Testis
Adipose_Subcutaneous
Adrenal_Gland
Breast_Mammary_Tissue
Minor_Salivary_Gland
Adipose_Visceral_Omentum
Whole_Blood
Brain_Cortex
Liver
Lung
Brain_Frontal_Cortex_BA9
Kidney_Cortex
rain_Anterior_cingulate_cortex_BA24
in_Nucleus_accumbens_basal_ganglia
Brain_Caudate_basal_ganglia
Brain_Putamen_basal_ganglia
Brain_Amygdala
Brain_Hypothalamus
Brain_Hippocampus
Brain_Spinal_cord_cervical_c-1
Brain_Substantia_nigra

#### LVESV

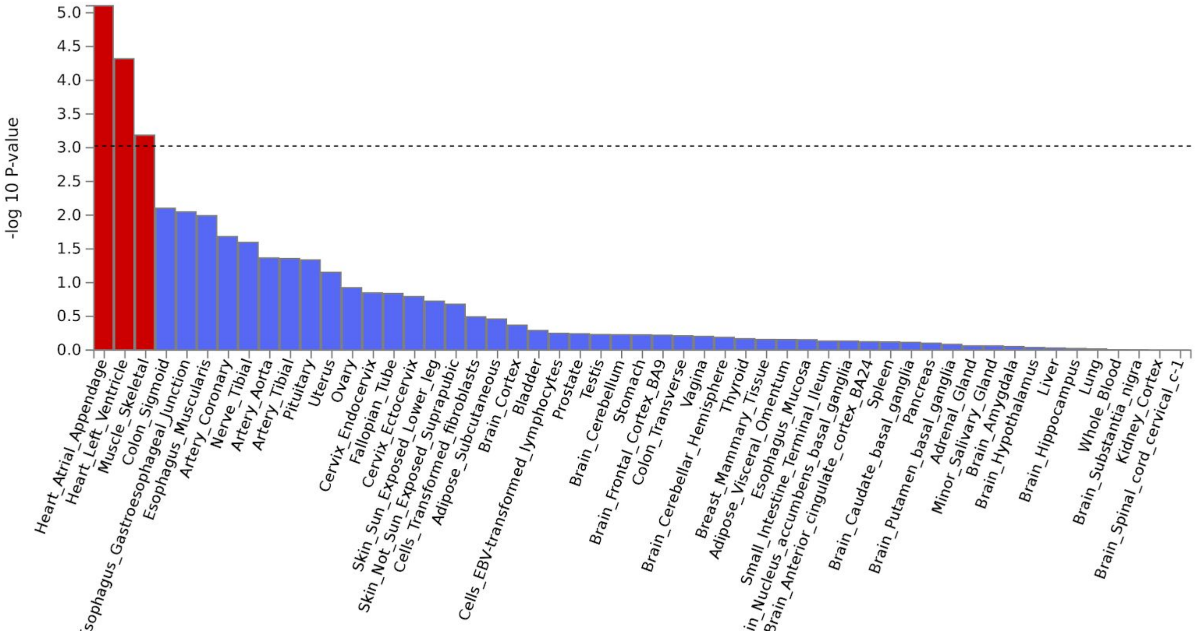
-log 10 P-value
0.0
0.5
1.0
1.5
2.0
2.5
3.0
3.5
4.0
4.5
5.0
Heart_Atrial_Appendage
Heart_Left_Ventricle
Muscle_Skeletal
Colon_Sigmoid
sophagus_Gastroesophageal_Junction
Esophagus_Muscularis
Artery_Coronary
Nerve_Tibial
Artery_Aorta
Artery_Tibial
Pituitary
Uterus
Ovary
Cervix_Endocervix
Fallopian_Tube
Cervix_Ectocervix
Skin_Sun_Exposed_Lower_leg
Skin_Not_Sun_Exposed_Suprapubic
Cells_Transformed_fibroblasts
Adipose_Subcutaneous
Brain_Cortex
Bladder
Cells_EBV-transformed_lymphocytes
Prostate
Testis
Brain_Cerebellum
Stomach
Brain_Frontal_Cortex_BA9
Colon_Transverse
Vagina
Brain_Cerebellar_Hemisphere
Thyroid
Breast_Mammary_Tissue
Adipose_Visceral_Omentum
Esophagus_Mucosa
Small_Intestine_Terminal_Ileum
in_Nucleus_accumbens_basal_ganglia
rain_Anterior_cingulate_cortex_BA24
Spleen
Brain_Caudate_basal_ganglia
Pancreas
Brain_Putamen_basal_ganglia
Adrenal_Gland
Minor_Salivary_Gland
Brain_Amygdala
Brain_Hypothalamus
Liver
Brain_Hippocampus
Lung
Whole_Blood
Brain_Substantia_nigra
Kidney_Cortex
Brain_Spinal_cord_cervical_c-1

LVEF

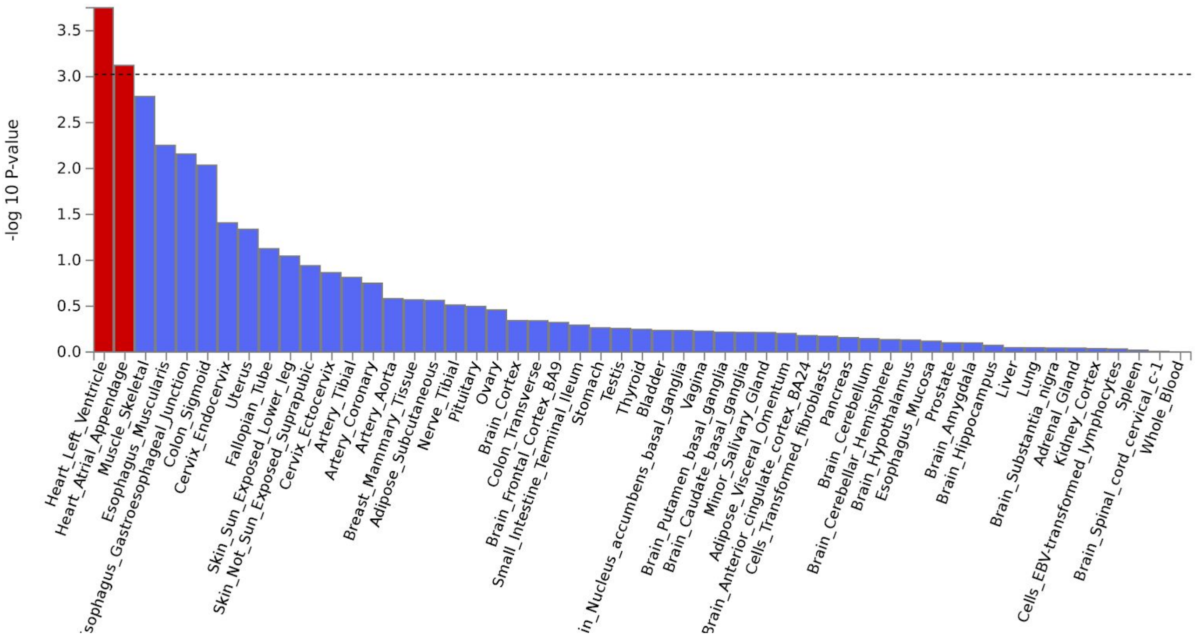
-log 10 P-value
0.0
0.5
1.0
1.5
2.0
2.5
3.0
3.5
Heart_Left_Ventricle
Heart_Atrial_Appendage
Muscle_Skeletal
Esophagus_Muscularis
sophagus_Gastroesophageal_Junction
Colon_Sigmoid
Cervix_Endocervix
Uterus
Fallopian_Tube
Skin_Sun_Exposed_Lower_leg
Skin_Not_Sun_Exposed_Suprapubic
Cervix_Ectocervix
Artery_Tibial
Artery_Coronary
Artery_Aorta
Breast_Mammary_Tissue
Adipose_Subcutaneous
Nerve_Tibial
Pituitary
Ovary
Brain_Cortex
Colon_Transverse
Brain_Frontal_Cortex_BA9
Small_Intestine_Terminal_Ileum
Stomach
Testis
Thyroid
Bladder
in_Nucleus_accumbens_basal_ganglia
Vagina
Brain_Putamen_basal_ganglia
Brain_Caudate_basal_ganglia
Minor_Salivary_Gland
Adipose_Visceral_Omentum
Brain_Anterior_cingulate_cortex_BA24
Cells_Transformed_fibroblasts
Pancreas
Brain_Cerebellum
Brain_Cerebellar_Hemisphere
Brain_Hypothalamus
Esophagus_Mucosa
Prostate
Brain_Amygdala
Brain_Hippocampus
Liver
Lung
Brain_Substantia_nigra
Adrenal_Gland
Kidney_Cortex
Cells_EBV-transformed_lymphocytes
Spleen
Brain_Spinal_cord_cervical_c-1
Whole_Blood

SV

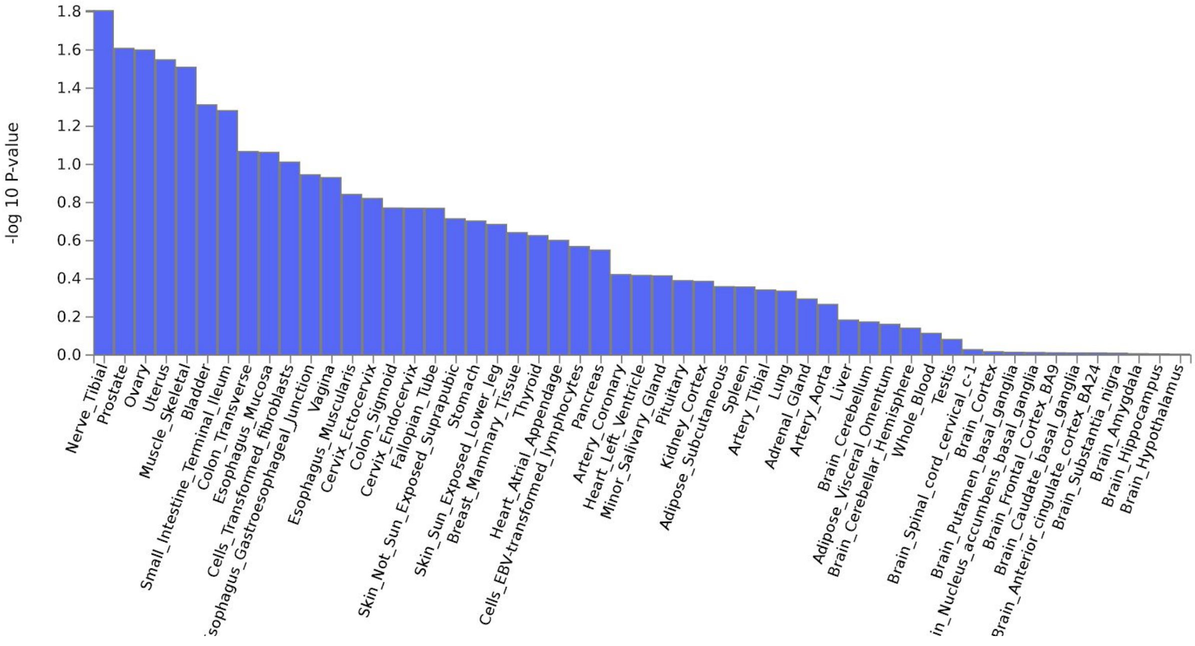
-log 10 P-value
0.0
0.2
0.4
0.6
0.8
1.0
1.2
1.4
1.6
1.8
Nerve_Tibial
Prostate
Ovary
Uterus
Muscle_Skeletal
Bladder
Small_Intestine_Terminal_Ileum
Colon_Transverse
Esophagus_Mucosa
Cells_Transformed_fibroblasts
sophagus_Gastroesophageal_Junction
Vagina
Esophagus_Muscularis
Cervix_Ectocervix
Colon_Sigmoid
Cervix_Endocervix
Fallopian_Tube
Skin_Not_Sun_Exposed_Suprapubic
Stomach
Skin_Sun_Exposed_Lower_leg
Breast_Mammary_Tissue
Thyroid
Heart_Atrial_Appendage
Cells_EBV-transformed_lymphocytes
Pancreas
Artery_Coronary
Heart_Left_Ventricle
Minor_Salivary_Gland
Pituitary
Kidney_Cortex
Adipose_Subcutaneous
Spleen
Artery_Tibial
Lung
Adrenal_Gland
Artery_Aorta
Liver
Brain_Cerebellum
Adipose_Visceral_Omentum
Brain_Cerebellar_Hemisphere
Whole_Blood
Testis
Brain_Spinal_cord_cervical_c-1
Brain_Cortex
Brain_Putamen_basal_ganglia
in_Nucleus_accumbens_basal_ganglia
Brain_Frontal_Cortex_BA9
Brain_Caudate_basal_ganglia
3rain_Anterior_cingulate_cortex_BA24
Brain_Substantia_nigra
Brain_Amygdala
Brain_Hippocampus
Brain_Hypothalamus

### LVEDVi

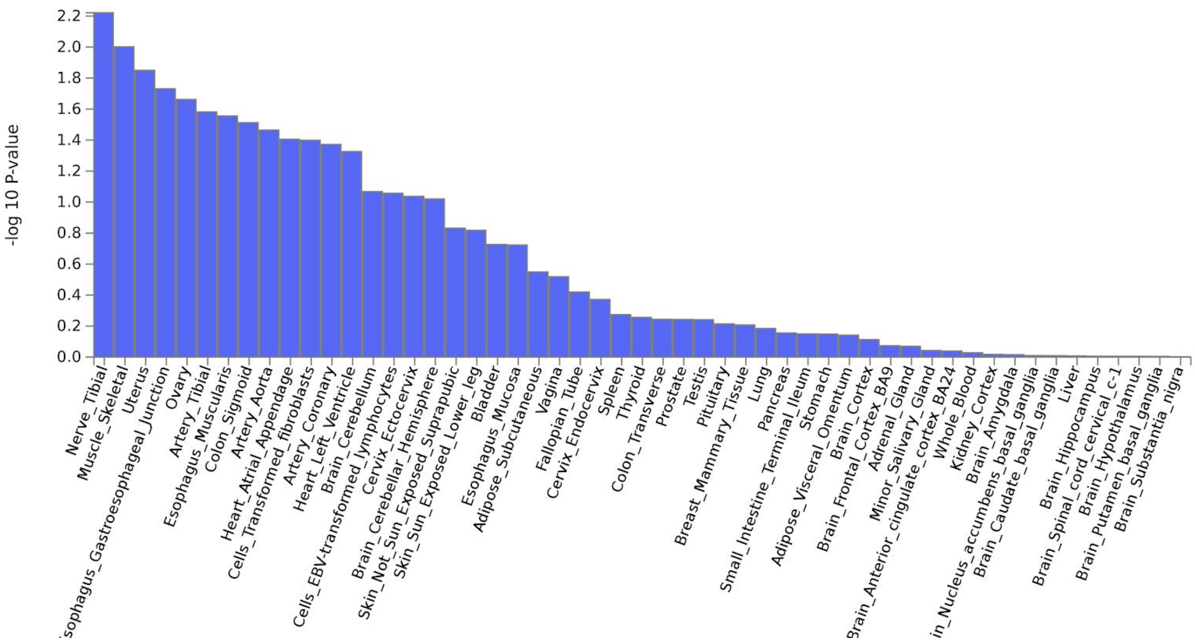


### LVESVi

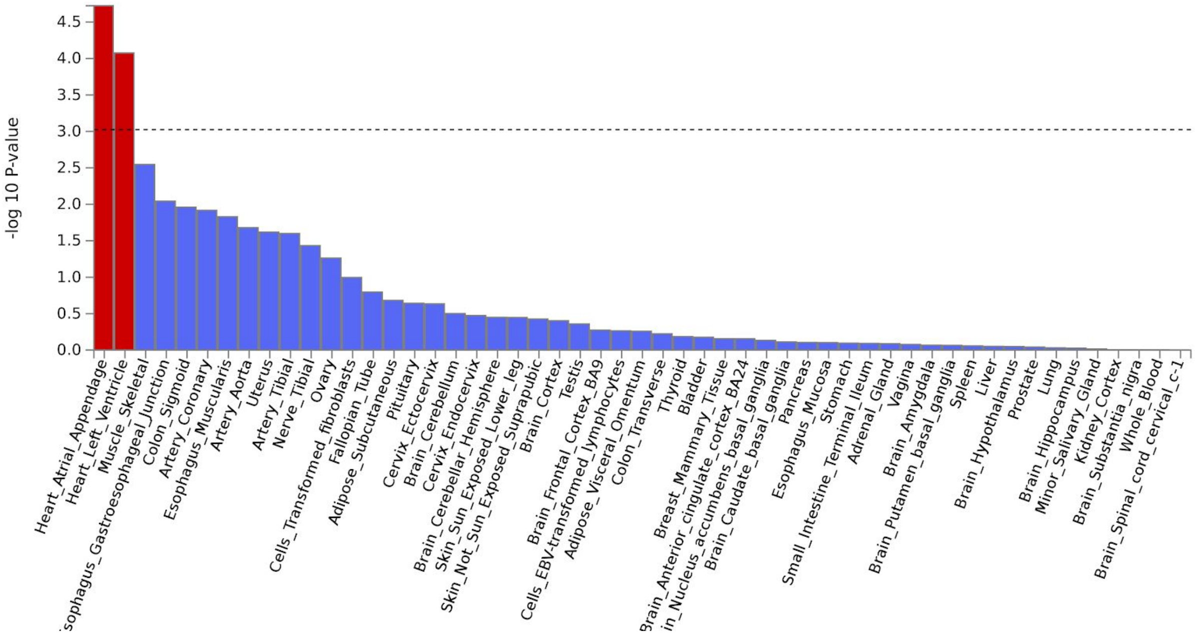

SVi

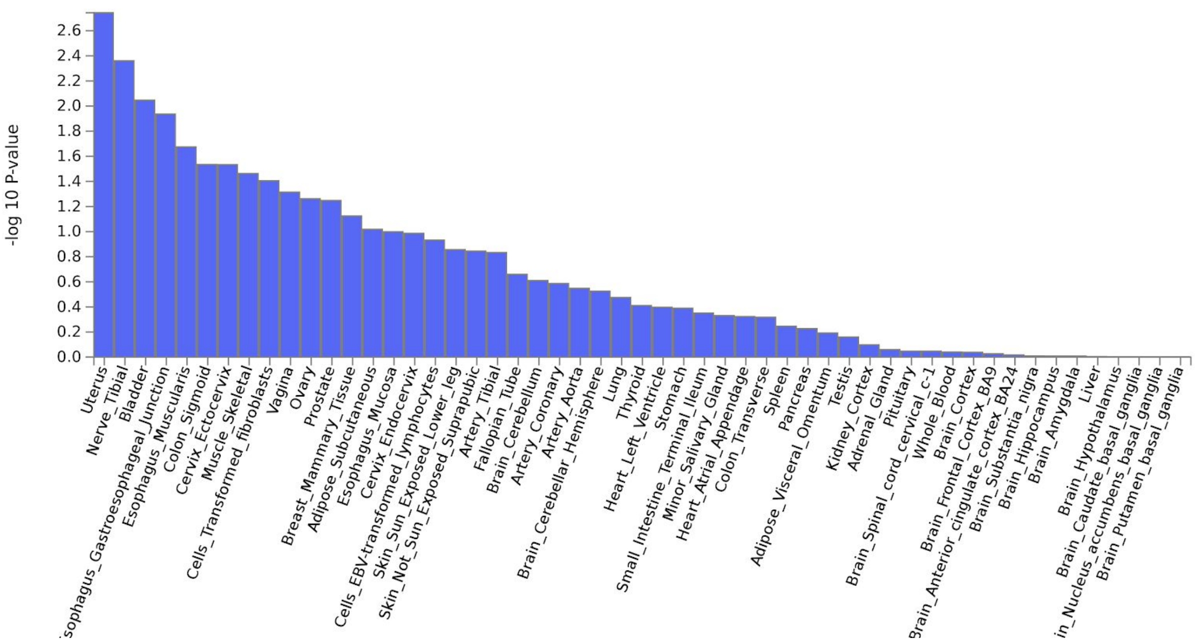

-log 10 P-value
0.0
0.2
0.4
0.6
0.8
1.0
1.2
1.4
1.6
1.8
2.0
2.2
2.4
2.6
Uterus
Nerve_Tibial
Bladder
Esophagus_Gastroesophageal_Junction
Esophagus_Muscularis
Colon_Sigmoid
Cervix_Ectocervix
Muscle_Skeletal
Cells_Transformed_fibroblasts
Vagina
Ovary
Prostate
Breast_Mammary_Tissue
Adipose_Subcutaneous
Esophagus_Mucosa
Cervix_Endocervix
Cells_EBV-transformed_lymphocytes
Skin_Sun_Exposed_Lower_leg
Skin_Not_Sun_Exposed_Suprapubic
Artery_Tibial
Fallopian_Tube
Brain_Cerebellum
Artery_Coronary
Artery_Aorta
Brain_Cerebellar_Hemisphere
Lung
Thyroid
Heart_Left_Ventricle
Stomach
Small_Intestine_Terminal_Ileum
Minor_Salivary_Gland
Heart_Atrial_Appendage
Colon_Transverse
Spleen
Pancreas
Adipose_Visceral_Omentum
Testis
Kidney_Cortex
Adrenal_Gland
Pituitary
Brain_Spinal_cord_cervical_c-1
Whole_Blood
Brain_Cortex
Brain_Frontal_Cortex_BA9
Brain_Anterior_cingulate_cortex_BA24
Brain_Substantia_nigra
Brain_Hippocampus
Brain_Amygdala
Liver
Brain_Hypothalamus
Brain_Caudate_basal_ganglia
Brain_Nucleus_accumbens_basal_ganglia
Brain_Putamen_basal_ganglia

### Supplemental Figure 6: GWAS loci are found nearby to more cardiomyopathy-related genes than expected by chance

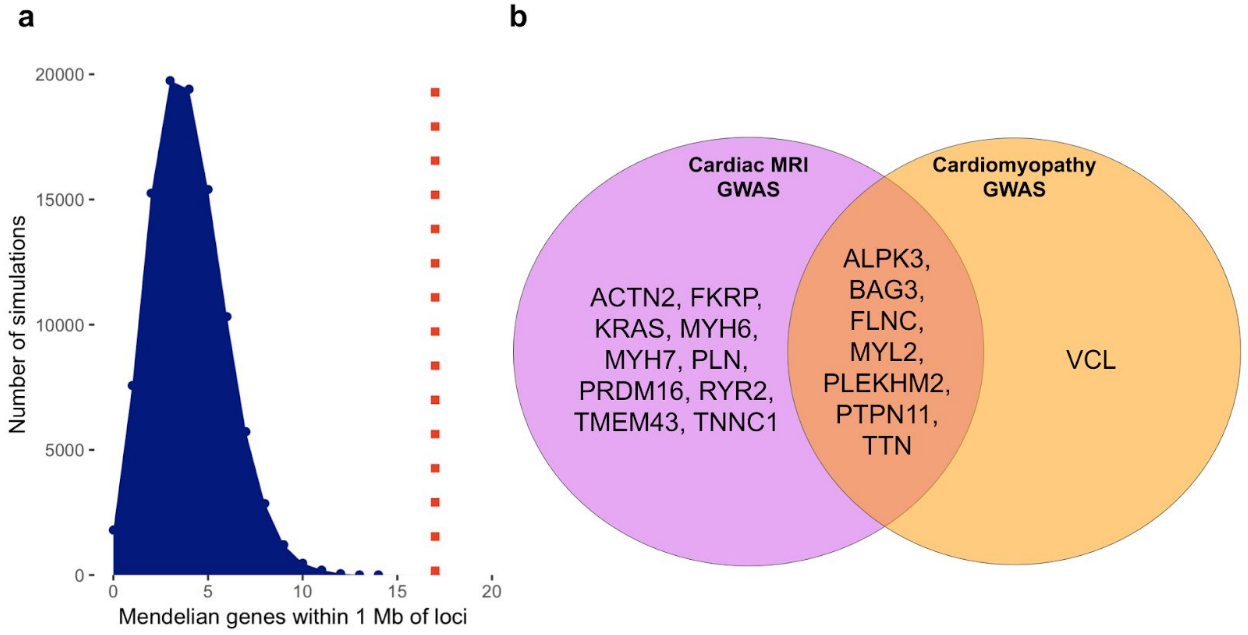


**a.** In 100,000 simulations of SNPs matched to our GWAS lead SNPs in their statistical properties, on average 4 Mendelian cardiomyopathy genes were located within 1 Mb of those SNPs (range: 0-14 genes). In contrast, 17 Mendelian genes were found within 1 Mb of the actual SNPs from our study (dotted red vertical line; $P < 1 \times 10^{-5}$). **b.** Mendelian cardiomyopathy-related genes found near genome-wide significant loci associated with cardiomyopathy and cardiac MRI phenotypes are plotted in a Venn diagram. Genes found only near loci discovered in this GWAS of cardiac MRI phenotypes are located in the pink circle (10 genes); those found only in prior cardiomyopathy GWAS appear in the yellow circle (1 gene); and those found in both are in the overlapping orange area (7 genes). In order to appear in this figure, a cardiomyopathy-related gene symbol from **Supplemental Table 4** had to fall within 1 Mb of a lead SNP from a GWAS.

### Supplemental Figure 7: PheWAS

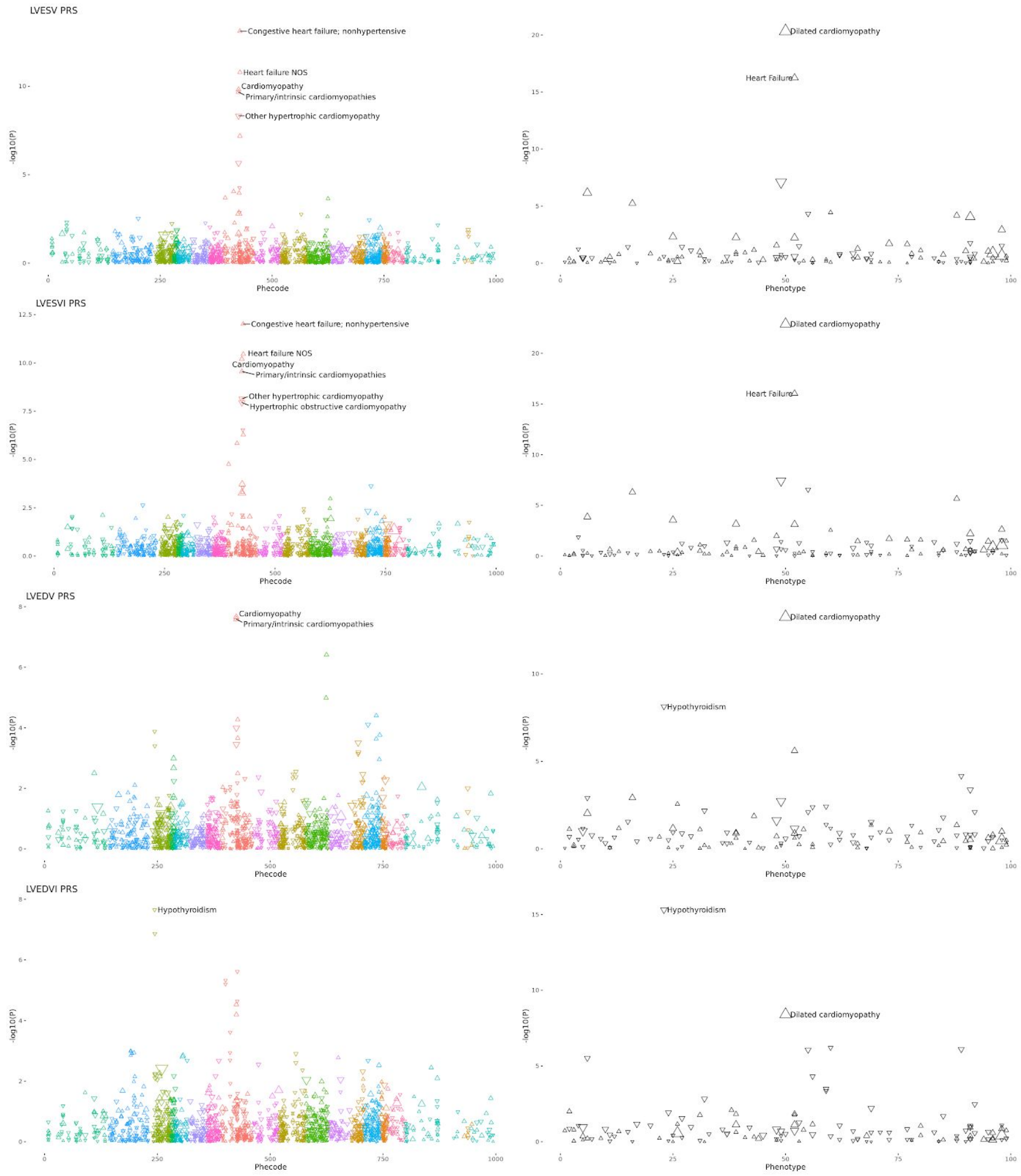

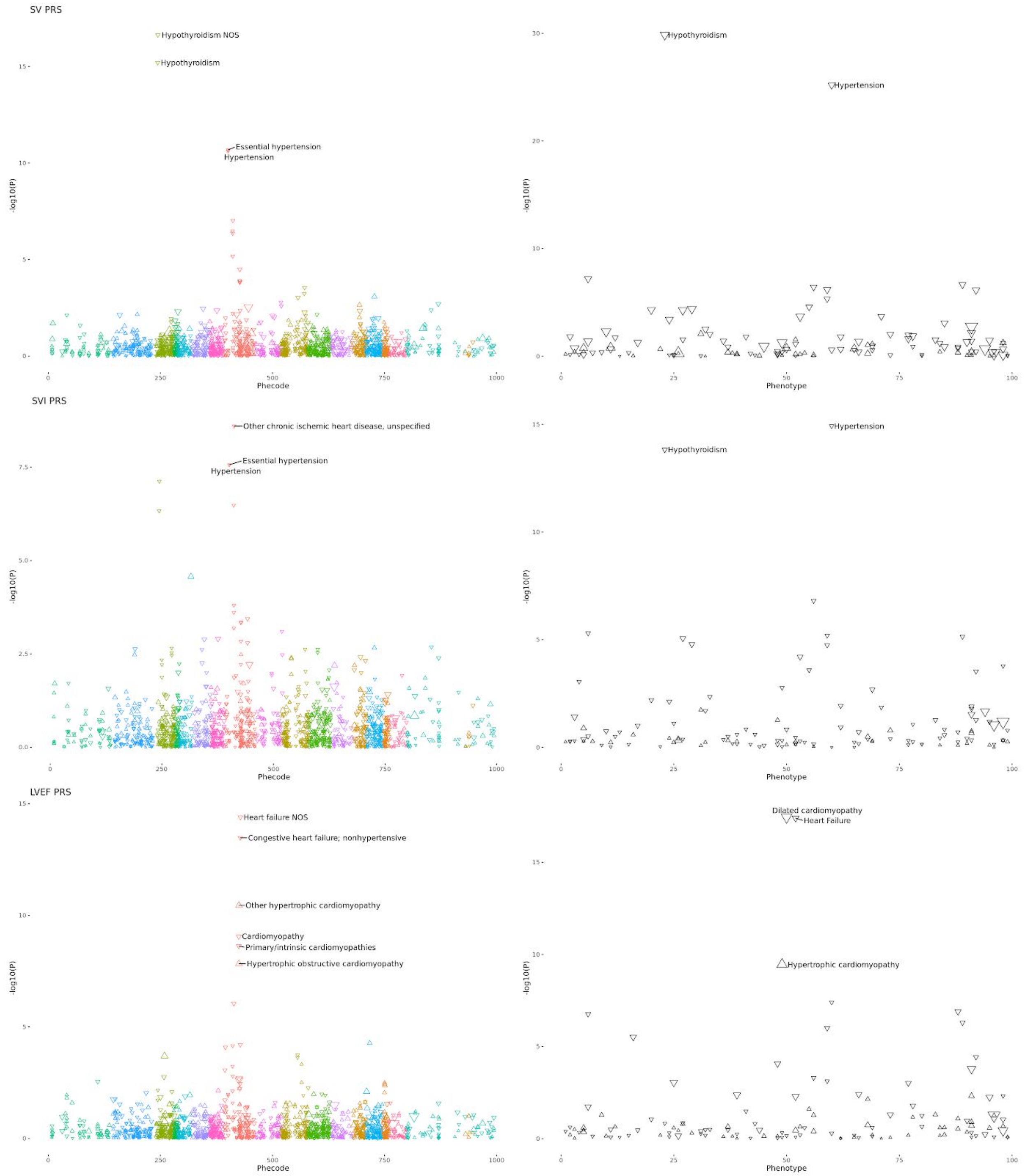


For each of the 7 traits for which a genome-wide association study was performed, the polygenic score was applied to two sets of phenotypes. On the left, colored by Phecode grouping, are Phecode-defined phenotypes. On the right in black-and-white are the curated phenotypes defined in **Supplemental Table 1**. The X-axis is manually defined and arbitrary, though attempts to cluster similar phenotypes. The Y-axis represents the -log10 of the P-value of association between the polygenic score and the phenotype in a logistic model adjusted for age at enrollment, the genotyping array, sex, and the first five principal components of ancestry. The most strongly associated phenotypes for each group are labeled, as are any phenotypes with association $P < 1 \times 10^{-20}$.
